# Multiple-labeled antibodies behave like single emitters in photoswitching buffer

**DOI:** 10.1101/2020.07.23.217125

**Authors:** Dominic A. Helmerich, Gerti Beliu, Markus Sauer

**Affiliations:** Department of Biotechnology and Biophysics, Biocenter, and Rudolf Virchow Center for Integrative and Translational BioImaging, University of Würzburg, 97074 Würzburg, Germany

## Abstract

The degree of labeling (DOL) of antibodies has so far been optimized for high brightness and specific and efficient binding. The influence of the DOL on the blinking performance of antibodies used in *direct* stochastic optical reconstruction microscopy (*d*STORM) has so far attained limited attention. Here, we investigated the spectroscopic characteristics of IgG antibodies labeled at DOLs of 1.1- 8.3 with Alexa Fluor 647 (Al647) at the ensemble and single-molecule level. Multiple-Al647-labeled antibodies showed weak and strong quenching interactions in aqueous buffer but could all be used for *d*STORM imaging with spatial resolutions of ∼ 20 nm independent of the DOL. Photon antibunching experiments in aqueous buffer demonstrate that the emission of multiple-Al647-labeled antibodies switches from classical to non-classical light in photoswitching buffer. We developed a model that explains the observed blinking of multiple-labeled antibodies and can be used advantageously to develop improved fluorescent probes for *d*STORM experiments.

## Introduction

The unique combination of molecule-specific labeling, minimal perturbation and three-dimensional (3D) imaging makes fluorescence microscopy the most popular imaging modality in cell biology. With the advent of single-molecule and super-resolution microscopy methods, we can now achieve a spatial resolution that is well below the diffraction limit of light microscopy, enabling invaluable insights into the spatial organization of cells and biological samples^1,2^. High-end fluorescence microscopy enables the quantitative study of the distribution, interactions and dynamics of proteins, molecular machines and other cellular components at the single-molecule level^3,4^. However, specific labeling of the molecule of interest with a fluorescent probe that shows a high fluorescence intensity and photostability is an absolute prerequisite. Usually, organic dyes^5-7^ and fluorescent proteins (FPs)^8^ are used for single-molecule fluorescence and super-resolution microscopy experiments. To minimize photobleaching especially in single-molecule tracking experiments semiconductor quantum dots (QDs) and fluorescent nanoparticles have been developed^9-11^. Furthermore, multiple dye-labeled probes such DNA-based FluoroCubes^12^, bottlebrush polymers with DNA-tipped bristles^13^ and biotin-avidin signal amplifications^14^ have been introduced in combination with immunolabeling and chemical tags to reduce photobleaching in highly sensitive fluorescence imaging experiments.

Commonly, a compromise has to be accepted when designing multiple dye-labeled probes. On the one hand, the overall size of the probe should be small and easy to functionalize to ensure efficient high density labeling of the molecule of interest. On the other hand, intermolecular quenching interactions between fluorophores, e.g. via formation of nonfluorescent dye aggregates should be avoided^12,13,15-17^. Generally, standard immunlabeling with a primary and secondary antibody is used for imaging of endogenous proteins. Since a typical IgG antibody carries numerous lysine residues it can be labeled with multiple dyes^18^ and represents thus a multiple dye-labeled probe. However, the degree of labeling (DOL) has to be kept below 6-8 dyes per antibody to minimize quenching effects and retain a high binding specificity and affinity^19,20^. Usually antibodies are optimized for high brightness and efficient binding to their corresponding epitopes, which typically results in degrees of labeling (DOL) of 2-5^16,20^.

Interestingly, the DOL of antibodies used in single-molecule localization microscopy experiments has so far been largely ignored. For example, single-molecule localization microscopy by *d*STORM^21^ requires multiple-labeled antibodies that show synchronized blinking in order to enable precise localization of spatially isolated probes and artifact-free image reconstruction. On closer inspection of dye-labeled antibodies used for immunostaining in previous *d*STORM experiments it becomes evident that most of them exhibited in fact a DOL ≥ 1^21-24^ implying that multiple-labeled antibodies show ‘collective’ blinking in photoswitching buffer. Synchronous or ‘collective’ blinking has been observed for multichromophoric systems where multiple fluorophores are positioned at nanometer distances and interact via various energy transfer processes including homo-FRET, singlet-triplet-FRET and other annihilation processes^25,26^. Interestingly enough some multichromophoric systems showed photon-antibunching, which is a characteristic signature of a single emitter^17,25-29^. Hence, multiple-labeled antibodies might in fact be possibly suitable for *d*STORM.

Here, we investigated the fluorescence characteristics of antibodies labeled with different DOLs and their performance in *d*STORM imaging. We show that multiple Alexa Fluor 647 (Al647) labeled IgG antibodies exhibit strong quenching interactions with increasing DOL but unaltered blinking performance in photoswitching buffer. Photon antibunching experiments in photoswitching buffer demonstrate that multiple-Al647-labeled IgG antibodies behave like single emitters in *d*STORM experiments.

## Results

### Fluorescence quenching in multiple-labeled antibodies

To investigate fluorescence quenching in multiple-labeled antibodies we conjugated the N-hydroxysuccinimidyl esters of carbocyanine dyes (Cy5 and Al647) to secondary goat anti-rabbit IgG antibodies via primary amines according to the manufacturer’s specifications. Using different dye-to-protein ratios, the resulting DOL varied between 1.1 and 8.3 as determined by absorption spectroscopy. Considering the size of an IgG antibody of ∼10 nm^30^ it becomes apparent that the fluorophores can interact in different ways. If the fluorophores are arranged so closely (< 2 nm) that their wave functions mix a new delocalized state can be produced as, for example in the B850 ring of the light harvesting complex 2 (LH2)^31^. Strong attractive interactions between fluorophores can also lead to stable complexes that change the excited electronic energy landscape and induce formation of fluorescent J- or non-fluorescent H-dimers^15,32,33^. On the other hand, if the fluorophores are spaced further they show weak electronic interactions, which can be well-described within the framework of the Förster theory^34,35^. In this weak coupling regime, electronic excitation transfer involves coupling of the transition moments of the fluorophores via Coulombic interactions, and the absorption spectrum constitutes the sum of the individual components.

To investigate Al647 interactions we performed absorption and fluorescence spectroscopy measurements of multiple-labeled IgG antibodies with varying DOL (**Fig. 1a** and **Supplementary Table 1**). Please note that an ensemble solution of an antibody with a certain DOL contains a mixture of antibody species with different labeling degree whose distribution is usually assumed to be Poissonian^20^. Al647 is structurally identical to the carbocyanine dye Cy5 but exhibits four instead of two sulfonate groups (**Supplementary Fig. 1**)^36^ and thus a more hydrophilic character than Cy5^37^. The absorption spectrum of Al647 in phosphate buffered saline (PBS), pH 7.4 shows the expected absorption maximum at 649 nm with a characteristics shoulder at ∼610 nm (**Fig. 1a**). Upon conjugation to antibodies the absorption spectra display an increase in absorption at 610 nm with increasing DOL indicating dimer formation between Al647 molecules^38,39^. Multiple-Cy5-labeled antibodies exhibit a more pronounced dimer-shoulder at ∼600 nm and accordingly also a lower fluorescence intensity indicating stronger fluorophore interactions (**Supplementary Fig. 2**). In the following we focused our investigations solely on multiple-Al647-labeled antibodies with DOLs of 1.1, 2.1, 4.1, and 8.3.

**Figure 1.**
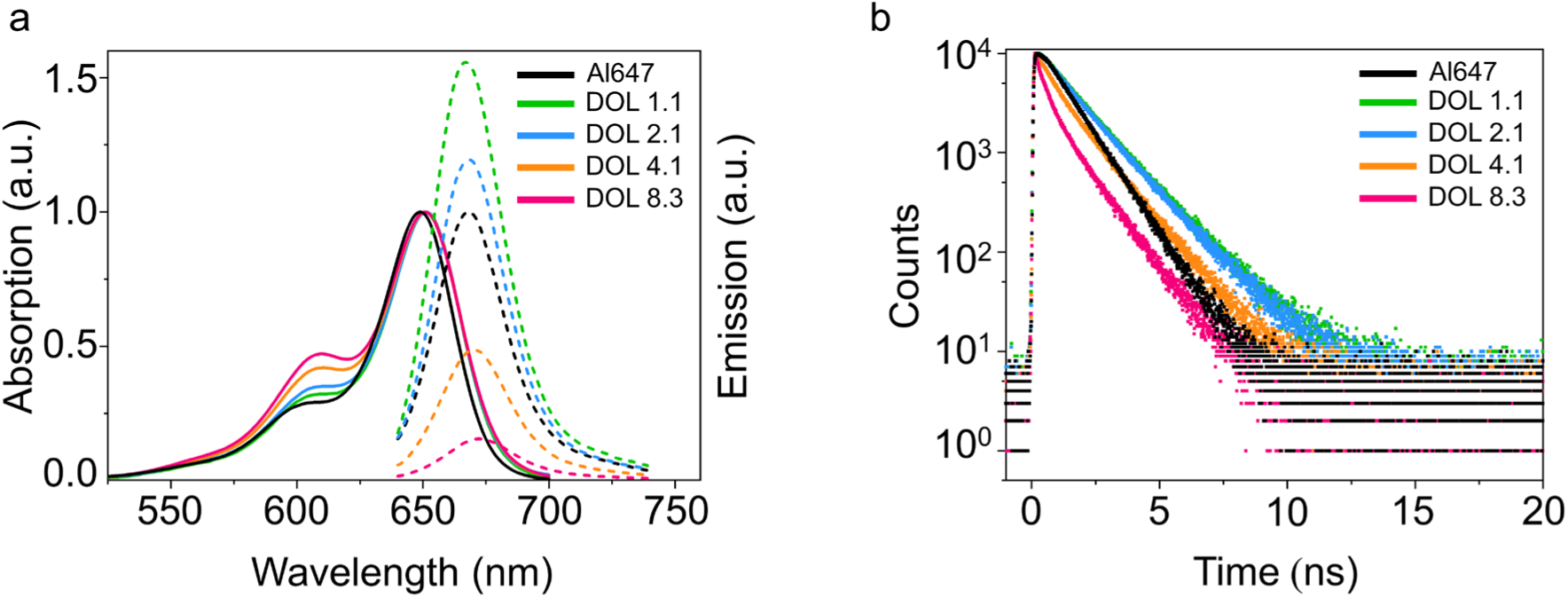
Ensemble absorption and fluorescence emission characteristics of multiple-Al647-labeled antibodies. (**a**) Normalized absorption and relative fluorescence emission spectra of multiple-Al647-labeled IgG antibodies measured in PBS, pH 7.4. (**b**) Fluorescence decay curves of multiple-Al647-labeled IgG antibodies measured in PBS, pH 7.4 at concentrations of ≤ 1 µM.

The absorption spectra of Al647 labeled antibodies exhibit a small bathochromic shift of the absorption maximum from 649 to 651 nm (**Fig. 1a**). The characteristic absorption spectra are independent of the antibody concentration demonstrating that the observed changes are not caused by antibody interactions, e.g. aggregation (**Supplementary Fig. 3a**). The fluorescence emission maximum shifts also bathochromic from 667 nm for DOL 1.1 to 672 nm for DOL 8.3 (**Table 1**). The fluorescence quantum yield increases slightly for DOL 1.1 and 2.1 and decreases then strongly for higher DOLs of 4.1 and 8.3 compared to the free dye in aqueous buffer (**Fig. 1a**). The observed dimer absorption arises from interactions between two or more dye monomers attached to the same antibody. These interactions result in splitting of the singlet excited state of the dimer into a long-wavelength and a short wavelength band on each side of the main maximum of the dye monomer (**Fig. 1a**)^38,40^. The ratio of the intensity of the long-wavelength band to that of the short-wavelength band depends on the relative position of interacting dye monomers. The fact that the short-wavelength band, i.e. the absorption shoulder at ∼610 nm dominates the absorption spectrum demonstrates that the Al647 molecules are nearly parallel aligned forming non-fluorescent H-dimers^15,32,33,38,40^. Ensemble absorption and fluorescence emission spectra of multiple-Al647-labeled antibodies in photoswitching buffer containing 100 mM mercaptoethylamine (MEA) showed identical spectra but a weak increase in fluorescence intensity in the presence of MEA (**Supplementary Fig. 3b**).

**Table 1.** On-state fluorescence intensity (I_ON_) in photons ms^-1^, on-state lifetime (τ_ON_) in milliseconds, off-state lifetime (τ_OFF_) in seconds, and average *xy* localization precision (Loc)^45^ in nanometers determined from immobilized individual multiple-Al647-labeled antibodies. Data were recorded in PBS, pH 7.6 containing 100 mM MEA in the absence and presence of oxygen scavenger. Samples were excited with circular polarized light at 640 with an irradiation intensity of 2 kW cm^-2^ without or with additional 405 nm light with 8 W cm^-2^ (+UV). Detection was performed at a frame rate of 125 Hz. Subsequent blink events of the same antibody were histogrammed and the average parameters determined. The average parameters of each antibody were then histogrammed to determine the mean values ± standard deviation (s.d.)

| | DOL | $I_{ON}$ ($ms^{-1}$) | $\tau_{ON}$ (ms) | $\tau_{OFF}$ (s) | Loc (nm) |
| --- | --- | --- | --- | --- | --- |
| MEA | 1.1 | $396.8 \pm 151.5$ | $10.2 \pm 1.7$ | $12.2 \pm 11.8$ | $5.1 \pm 1.9$ |
| | 2.1 | $398.5 \pm 156.7$ | $10.2 \pm 2.1$ | $10.1 \pm 9.2$ | $5.2 \pm 1.9$ |
| | 4.1 | $396.0 \pm 160.0$ | $10.2 \pm 1.9$ | $11.8 \pm 11.2$ | $5.2 \pm 1.9$ |
| | 8.3 | $394.2 \pm 159.6$ | $10.3 \pm 2.1$ | $11.2 \pm 11.2$ | $5.8 \pm 2.1$ |
| MEA, oxygen scavenger | 1.1 | $356.3 \pm 132.3$ | $11.0 \pm 2.1$ | $21.5 \pm 18.7$ | $5.1 \pm 1.7$ |
| | 2.1 | $418.1 \pm 161.6$ | $10.9 \pm 2.2$ | $22.4 \pm 19.2$ | $4.7 \pm 1.6$ |
| | 4.1 | $398.2 \pm 154.7$ | $10.9 \pm 2.3$ | $25.9 \pm 23.8$ | $4.8 \pm 5.1$ |
| | 8.3 | $389.9 \pm 154.5$ | $10.9 \pm 2.5$ | $24.4 \pm 22.4$ | $5.1 \pm 1.9$ |
| MEA, oxygen scavenger, +UV | 1.1 | $367.7 \pm 181.5$ | $11.0 \pm 2.8$ | $12.2 \pm 22.6$ | $5.3 \pm 1.8$ |
| | 2.1 | $369.0 \pm 171.0$ | $10.8 \pm 2.4$ | $13.6 \pm 25.3$ | $5.3 \pm 1.9$ |
| | 4.1 | $357.8 \pm 177.0$ | $11.1 \pm 3.3$ | $10.8 \pm 19.8$ | $5.5 \pm 1.9$ |
| | 8.3 | $367.8 \pm 167.4$ | $11.1 \pm 3.3$ | $13.7 \pm 24.9$ | $5.3 \pm 1.9$ |

For free Al647 we measured a monoexponential fluorescence lifetime of 1.09 ns (**Supplementary Fig. 1c**). Upon conjugation to the IgG antibody the fluorescence decays become biexponential and the average lifetimes increase for the two lower DOLs of 1.1 and 2.1 as already reported for conjugation of Al647 to proteins (**Supplementary Table 1**)^37^. The occurrence of multiexponential decays upon conjugation to proteins can be explained by changes in the cis/trans isomerization possibilities of carbocyanine dyes^41^. For the higher DOLs of 4.1 and 8.3, we measured a strong decrease in the average fluorescence lifetime reflecting the strong quenching observed in steady-state emission spectra (**Figs. 1a**,**b** and **Supplementary Table 1**). Overall, our ensemble data demonstrate strong fluorophore interactions in multiple Al647-labeled antibodies.

### Fluorescence lifetime imaging microscopy (FLIM) with multiple-Al647-labeled antibodies

Since the fluorescence decay times of multiple-Al647-labeled antibodies are controlled by the DOL (**Fig. 1b** and **Supplementary Table 1**) we tested if they can be used for fluorescence lifetime imaging microscopy (FLIM)^42,43^. To enable specific immunostaining of two different cellular proteins with secondary IgG antibodies, we labeled a secondary goat anti-mouse IgG with Al647 with a DOL of 0.9. The ensemble average fluorescence lifetimes of the goat anti-rabbit IgG with DOL 8.3 and goat anti-mouse IgG with DOL 0.9 were determined to 0.38 ns (**Supplementary Table 1**) and 1.49 ns (**Fig. 2a**), respectively, and should thus be easily distinguishable in FLIM experiments. Next, we immunostained clathrin with secondary goat anti-rabbit IgG DOL 8.3 and ß-tubulin with secondary goat anti-mouse IgG DOL 0.9 in COS7 cells. FLIM images of cells stained either with DOL 0.9 or DOL 8.3 (**Figs. 2b,c**) demonstrate that the average lifetimes of the two antibodies enable their unequivocal identification. Composite FLIM images of cells labeled with the two antibodies display that clathrin-coated pits and tubulin filaments can be unequivocally identified based on the different fluorescence lifetimes of multiple-Al647-labeled antibodies (**Fig. 2d**). A closer look at the FLIM images reveals that some of the DOL 8.3 antibodies label also other intracellular structures, which corroborates previous studies showing that multiple labeled antibodies exhibit a decreased affinity^20^. Interestingly, these unspecific binding antibodies exhibit a longer fluorescence lifetime, which can seriously complicate the interpretation of FLIM imaging experiments. (**Fig. 2b**). Nevertheless, our data clearly evidence the use of multiple-Al647-labeled antibodies with different DOL for ‘multi-target’ FLIM^43^.

**Figure 2.**
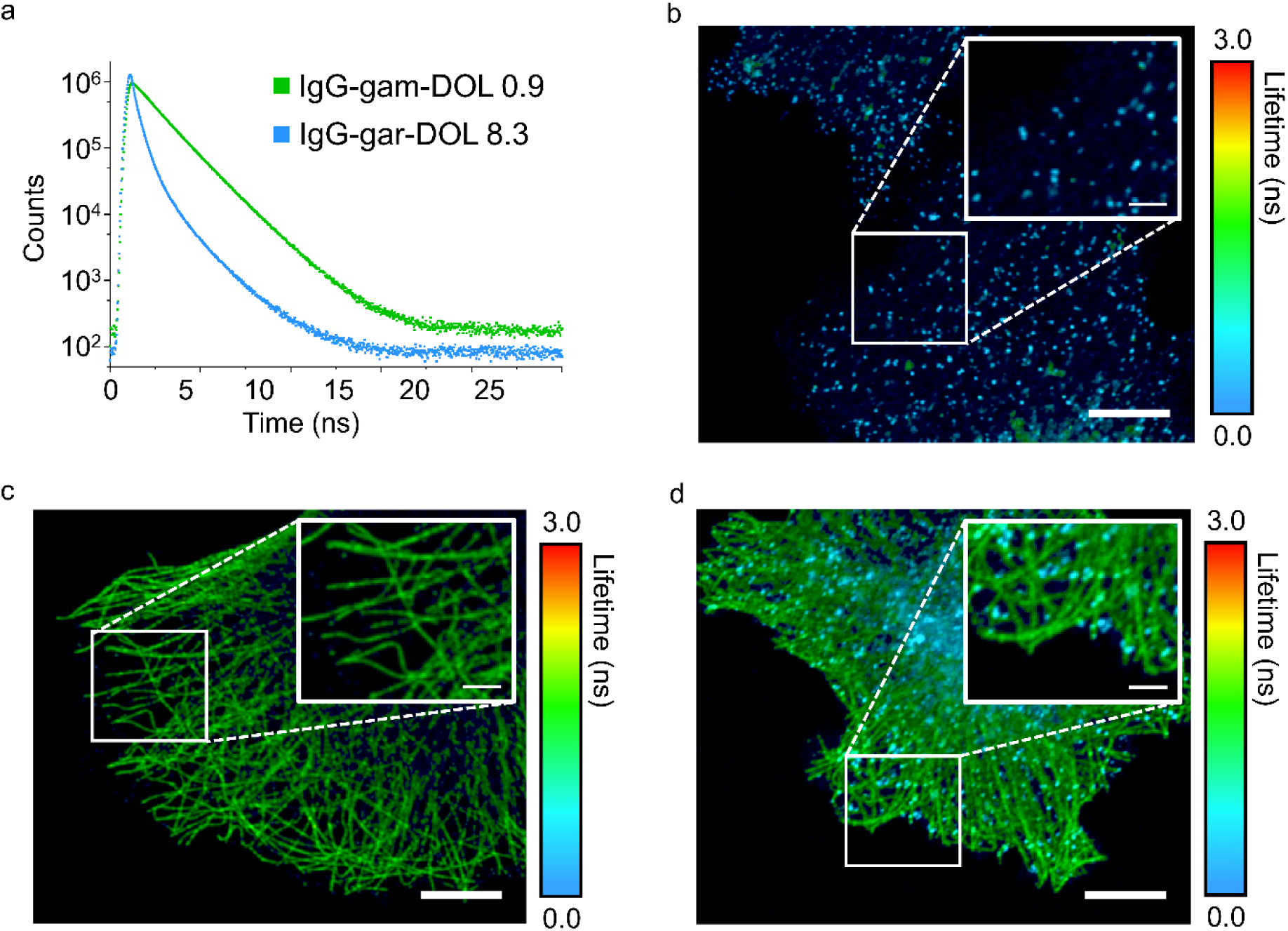
FLIM with multiple-AL647-labeled antibodies. (**a**) Fluorescence decays of goat anti-rabbit IgG DOL 8.3 and goat anti-mouse IgG DOL 0.9 excited at 640 nm. The decays were recorded at the emission maxima in PBS, pH 7.4. (**b**) FLIM image of a COS7 cell immunostained with mouse anti-clathrin primary and goat anti-mouse secondary antibodies with DOL 8.3. (**c**) FLIM image of a COS7 cell immunostained with rat anti-ß-tubulin primary and goat anti-rabbit secondary antibodies with DOL 0.9. (**d**) FLIM composite image of a COS7 cell immunostained with rat anti-ß-tubulin primary and mouse anti-clathrin primary and corresponding secondary multiple-Al647-labeled antibodies with different DOL. Scale bars, 10 µm.

### *d*STORM with multiple-Al647-labeled antibodies

Next, we investigated the performance of multiple-Al647-labeled antibodies in *d*STORM experiments and immunostained microtubules in COS7 cells. Using a photoswitching buffer containing 100 mM mercaptoethylamine (MEA) with and without oxygen scavenger and additional UV activation at 405 nm all antibodies showed distinct blinking independent of the DOL and the polarization of the irradiation laser at 641 nm, and enabled reconstruction of high quality *d*STORM images (**Figure 3, Supplementary Videos 1-4** and **Supplementary Figs. 4-7**). Particularly noteworthy here is the observed strong unspecific background labeling of higher DOL antibodies, which has also been seen in our FLIM experiments (**Fig. 2b**) corroborating again the finding that multiple-labeled antibodies exhibit a decreased binding affinity^20^. To investigate the blinking performance of multiple Al647-labeled antibodies in more detail we determined the average on- and off-times per blinking event as well as the fluorescence intensities (photons ms^-1^) in the on-state under different experimental conditions (**Supplementary Table 2**). Experiments under linear and circular polarized excitation revealed high quality *d*STORM images with similar on-state fluorescence intensities (**Supplementary Figs. 4-7** and **Supplementary Table 2**) and thus confirmed that Al647-labeled antibodies exhibit a high rotational mobility^44^ and can be used advantageously for *d*STORM imaging.

**Figure 3.**
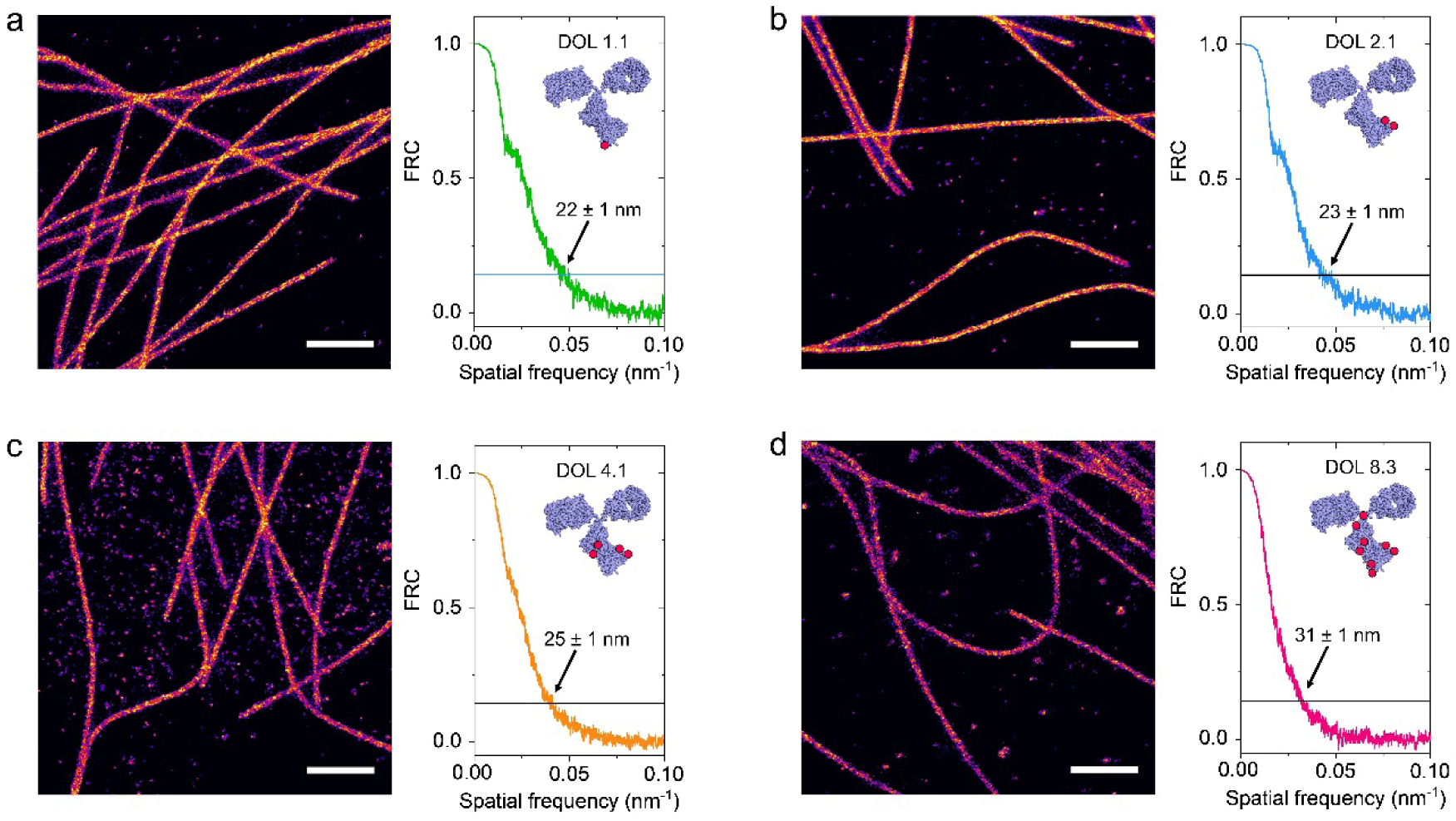
*d*STORM images of microtubules labeled with secondary multiple-Al647-labeled antibodies. *d*STORM images of COS7 cells immunostained with (**a**) DOL 1.1, (**b**) DOL 2.1, (**c**) DOL 4.1 and (**d**) DOL 8.3. Fourier ring correlation (FRC) analysis^46^ estimated spatial resolutions of 22 ± 1 nm (DOL 1.1), 23 ± 1 nm (DOL 2.1), 25 ± 1 nm (DOL 41.), and 31 ± 1 nm (DOL 8.3). *d*STORM images were recorded using circular polarized laser excitation at 641 nm with an irradiation intensity of 2 kW cm^-2^. Buffer conditions: 100 mM MEA with oxygen scavenger, pH 7.6. Scale bars, 1 µm.

Surprisingly, the on-state fluorescence intensities of single multiple-labeled antibodies with different DOLs are nearly identical in photoswitching buffer (**Supplementary Table 2**). This is in strong contrast to the ensemble fluorescence intensities measured in PBS buffer, which decreased strongly with increasing DOL (**Fig. 1a**). The average on-times of multiple-Al647-labeled antibodies were also unaffected by the DOL and the presence of oxygen scavenger, only the off-state lifetimes increased and decreased slightly in the presence of oxygen scavenger and additional irradiation at 405 nm, respectively (**Supplementary Table 2**). Accordingly, we determined similar localization precisions for the different DOLs of ∼ 6 nm (**Supplementary Table 2**)^45^. In addition, Fourier ring correlation analysis^46^ of microtubule *d*STORM images was performed and provided spatial resolutions of ∼ 20 nm for the lower DOL antibodies (DOL 1.1 – 4.1) and a slightly worse resolution for the highest DOL antibody (DOL 8.3), which might be caused by nonspecific background labeling (**Figure 3, Supplementary Fig. 8 and Supplementary Table 2**).

To investigate the blinking performance of multiple-Al647-labeled antibodies at the single-molecule level in more detail unperturbed by high emitter density artifacts, e.g. at crossing microtubule filaments, we immobilized antibodies on a glass coverslip with a surface density appropriate for single-molecule studies. Widefield single-molecule imaging experiments in photoswitching buffer allowed us to determine the average on- and off-state lifetimes and on-state fluorescence intensities under identical conditions as used in the *d*STORM experiments (**Table 1**). The determined values are very similar to the values determined from *d*STORM experiments (**Supplementary Table 2**) but underline even clearer that all blinking parameters of multiple-Al647-labeled antibodies appear to be independent of the DOL in photoswitching buffer. We determined fluorescence intensities in the on-state of ∼ 400 photons ms^-1^ and average on-state lifetimes of ∼ 10 ms irrespective of the experimental conditions and DOL. However, the data show more clearly the expected increase in off-state lifetimes in the presence of oxygen scavenger from ∼ 11 s to ∼ 24 s and decrease upon additional irradiation at 405 nm to ∼ 13 ms of multiple-Al647-labeled antibodies in photoswitching buffer (**Table 1**).

Taking together, our results of ensemble and *d*STORM studies provide support for the conclusion that electronic interactions between Al647 fluorophores such as the formation of non-fluorescent H-dimers are strongly destabilized at the single-molecule level in photoswitching buffer. In order to understand the blinking of multiple-Al647-labeled antibodies we have to revisit the photoswitching mechanism in thiol buffer. It has been discovered that efficient photoconversion of carbocyanine dyes from the on-state to the non-fluorescent off-state requires the presence of millimolar concentrations of a thiol, e.g. MEA, a pH > 7.0 and irradiation intensities of ≥ 1 kW cm^-2^^21-23^. Furthermore, it is generally accepted that the fluorophore has to be pumped into the triplet state before it reacts with the thiolate to form a long-lived, reduced off-state which absorbs at shorter wavelengths^47,48^. The off-state is relatively stable in aqueous solutions at room temperature and can be transferred back to the singlet ground state upon oxidation with molecular oxygen, which is facilitated upon irradiation at 405 nm. As a consequence, the lifetime of the off-state is mainly controlled by the redox properties of the fluorophore, the intensity of UV irradiation, and the oxygen concentration of the solution^22,48^. Hence, the off-state lifetime of multiple-labeled antibodies increases in the presence of oxygen scavenger and decrease again upon additional irradiation at 405 nm (**Table 1**).

### Collective blinking of multiple-Al647-labeled antibodies in photoswitching buffer

The questions which remain to be answered are: why do multiple-labeled antibodies show collective blinking in photoswitching buffer and why don’t we see any photophysical difference in their blinking performance? Our ensemble data showed that conjugation of Al647 to antibodies facilitates H-dimer formation because each protein exhibits only a limited number of preferential reaction sites, which might lead to ‘local’ crowding of fluorophores even at low DOLs^16^. The presence of 100 mM MEA in the photoswitching buffer does not alter the absorption spectra and, therefore, not the monomer-dimer equilibrium of multiple-labeled antibodies in the electronic singlet ground state (**Supplementary Fig. 3b**). However, it has been shown that fluorescent monomers and non-fluorescent dimers coexist in an equilibrium in aqueous solvents^49,50^. Fluorescence correlation spectroscopy (FCS) experiments with multiple-Al674-labeled antibodies showed fluorescence fluctuations in the nanosecond to microsecond time range of the FCS-curve in addition to the commonly observed cis-trans isomerization of carbocyanine dyes (**Supplementary Fig. 9**)^41^. These data are indicative of a monomer-dimer equilibrium in the microsecond to nanosecond time range between individual Al647 fluorophores conjugated to the same antibody.

Therefore, upon excitation monomers will be removed from the monomer-dimer equilibrium and transferred to the triplet-state via intersystem crossing followed by reaction with the thiolate to generate the long-lived off-state^47,48^. Since the off-state lifetime is three orders of magnitude longer than the on-state lifetime (**Table 1** and **Supplementary Table 2**) and monomers are preferentially excited because of their higher extinction coefficient^51^, dye monomers will accumulate in the off-state until they are transferred back to the singlet ground-state. Since the on-state exhibits a much shorter lifetime its fluorescence intensity is dominated by Al647 monomers.

### Single-molecule fluorescence trajectories of multiple-Al647-labeled antibodies

To verify our hypothesis, we investigated the photophysical characteristics of multiple-Al647-labeled antibodies in more detail by confocal single-molecule fluorescence microscopy. After scanning of surfaces individual immobilized antibodies were selected from the image, positioned in the laser focus and their fluorescence trajectory recorded until irreversible photobleaching occurred. Dynamic fluctuations in the properties of the fluorescence from single chromophores can reveal a wealth of information regarding the molecular motions of the molecule itself and the environment in which it resides, as well as the nonradiative energy-transfer processes between the fluorophore and other chemical species in close proximity^15^. To extend the observation time (increase the photostability) of single carbocyanine dyes in the laser focus we removed oxygen and used trolox/troloxquinone as efficient ROXS buffer^52,53^. First we acquired fluorescence trajectories of a double-stranded DNA construct labeled with a single Al647 molecule. Single Al647 molecules showed constant fluorescence intensity with some intensity fluctuations before single-step photobleaching occurred, typically after a few seconds (**Supplementary Fig. 10a**). In strong contrast, single Cy5 molecules displayed blinking with short on-states and long off-states in photoswitching buffer (100 mM MEA and oxygen scavenger) (**Supplementary Fig. 10c**).

Multiple-Al647-labeled antibodies showed partially one- and two-step photobleaching in trolox buffer for low DOLs but also more complex fluorescence fluctuations for higher DOLs as expected from multichromophoric systems with dynamic quenching interactions between individual fluorophores (**Figs. 4a-d**). For example, while the majority of DOL 4.1 and 8.3 antibodies showed strong fluorescence intensity fluctuations with temporarily strong increases in fluorescence intensity (**Figs. 4c**,**d**), some displayed stepwise photobleaching with a gradual decrease in fluorescence intensity as expected for independently emitting fluorophores (**Supplementary Fig. 11a**). As recently observed for multiple-labeled DNA origami^12^, the total photon output and on-time as a measure of photostability increased with increasing DOL in trolox buffer. On the other hand, in photoswitching buffer containing 100 mM MEA and oxygen scavenger multiple-Al647-labeled antibodies displayed on/off switching independent of the DOL of the antibody (**Figs. 4f-i**).

**Figure 4.**
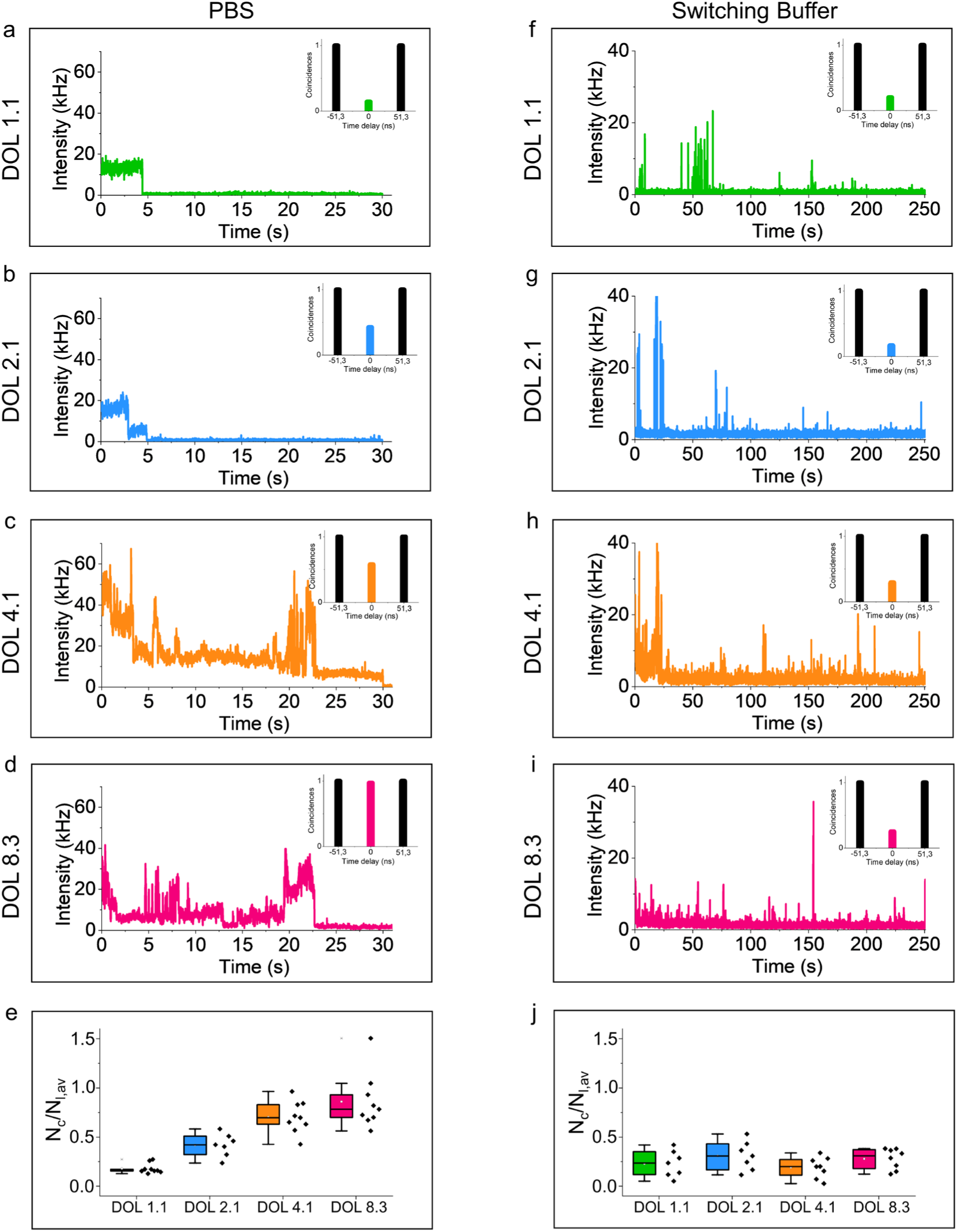
Single-molecule fluorescence intensity trajectories of multiple-AL647-labeled antibodies. (**a-d**) Fluorescence trajectories recorded for multiple-Al647-labeled antibodies with DOL 1.1 – 8.3 measured in PBS excited at 640 nm with 1 kW cm^-2^ (10 ms binning). The insets show the corresponding simplified interphoton time (coincidence) histograms containing the zero time delay and the first lateral peaks. (**e**) *N*_*c*_*/N*_*I,av*_ ratios measured for n = 7-9 single-molecule trajectories of multiple-Al647-labeled antibodies with DOL 1.1 – 8.3 in trolox buffer. (**f-i**) Fluorescence trajectories recorded for multiple-Al647-labeled antibodies with DOL 1.1 – 8.3 measured in photoswitching buffer containing 100 mM MEA and oxygen scavenger, excited at 640 nm with ∼1 kW cm^-2^ (10 ms binning). The insets show the corresponding simplified interphoton time (coincidence) histograms containing the zero time delay and the first lateral peaks. (**e**) *N*_*c*_*/N*_*I,av*_ ratios measured for n = 7-9 single-molecule trajectories of multiple-Al647-labeled antibodies with DOL 1.1 – 8.3 in photoswitching buffer.

### Photon antibunching of multiple-Al647-labeled antibodies in trolox buffer

The observation of transient dark states, i.e., when the fluorescence intensity reversibly drops to the background despite continuous illumination, is usually assumed to be an indication that the signal is from a single molecule, as multiple emitters would not be expected to go into these dark states simultaneously. To unequivocally demonstrate that multiple-Al647-labeled antibodies behave like a single emitter in photoswitching buffer we performed photon antibunching experiments. Photon antibunching experiments take advantage of the fact that the probability of emitting two consecutive photons drops to zero for a single emitter for time intervals shorter than the excited-state lifetime. After photon emission, a molecule must be re-excited and wait, on average, one fluorescence lifetime before another photon can be emitted. For sufficiently short laser pulses the number of photon-pairs detected per laser pulse can be used to determine whether the emission is from one or more independently emitting quantum systems. Therefore, we used the classical Hanbury-Twiss and Brown coincidence setup^54^ in combination with pulsed excitation to determine the number of independently emitting Al647 molecules per multiple-labeled antibody^25-29,55,56^. To record single-molecule fluorescence transients, single antibodies were selected from an image scan and positioned in the laser focus of a confocal microscope. After collection of the fluorescence by the same objective, the signal is separated by a 50/50 beam splitter and focused onto two avalanche photodiodes (APDs).

As expected for an ensemble of fluorophores, the distribution of interphoton times (coincidences) collected from a 200 nm fluorescent microsphere showed that the central peak is identical in intensity with the lateral ones (**Supplementary Fig. 12**). The central peak of the distribution at a delay time of 0 ns corresponds to photon pairs induced by the same laser pulse. In all other cases, interphoton times are distributed around a multiple of the repetition rate of the laser of 19.5 MHz, i.e. one peak every 51.3 ns. For a single emitter such as a double-stranded DNA construct labeled with a single Al647 molecule, the central peak decreases strongly since only one photon can be emitted after excitation (**Supplementary Figs. 10b**,**d**). Theoretically, one would expect that the central peak vanishes completely for a single emitter whose excited state lifetime is substantially longer than the laser pulse length. However, practically accidental coincidences due to background-background, signal-background, and background-signal contribute to the central peak (**Supplementary Fig. 13**). The coincidence histograms can be analyzed to determine the ratio of the number of photon pairs detected in the central peak, *N*_*c*_, at delay time zero, to the average number in the lateral peaks, *N*_*l,av*_. For *N*_*l,av*_, we used the average number of events in the nearest 8 peaks, 4 to each side of the zero delay time peak. For simplification, we show partially only the first lateral peaks (**Supplementary Figs. 10b**,**d**).

Since the intensity of the central peak contains information about the number of independently emitting molecules, the *N*_*c*_/*N*_*l,av*_ ratio can be used for molecular counting. For example, neglecting background, *N*_*c*_/*N*_*l,av*_ ratios of 0.0, 0.5, 0.67, and 0.75 are expected for 1-4 independently emitting fluorophores, respectively^25-29,55,56^. From the interphoton times histogram recorded from a single double-stranded DNA construct labeled with a single Al647 molecule we determined *N*_*c*_/*N*_*l,av*_ ratios of 0.03 ± 0.004 and 0.18 ± 0.04 in PBS and photoswitching buffer, respectively (**Supplementary Figs. 10b**,**d**).

Photon antibunching experiments of multiple-Al647-labeled antibodies in trolox buffer exhibited *N*_*c*_/*N*_*l,av*_ ratios largely expected for multiple independently emitting fluorophores (**Figs. 4a-d**). With increasing DOL the average *N*_*c*_/*N*_*l,av*_ ratio measured for several typical fluorescence trajectories increased as well from 0.18 ± 0.05 (DOL 1.1) via 0.42 ± 0.12 (DOL 2.1), and 0.70 ± 0.16 (DOL 4.1) to 0.86 ± 0.28 (DOL 8.3) matching the theoretical expectations for multichromophoric systems containing 1-8 dynamically interacting (quenched) fluorophores (**Fig. 4e**)^25-27,56^. As already mentioned, some higher DOL antibodies showed stepwise photobleaching (**Supplementary Fig. 11a**). Coincidence histograms generated for the consecutive bleaching steps demonstrate that the intact multiple-Al647-labeled antibody with DOL 4.1 exhibited a *N*_*c*_*/N*_*I,av*_ ratio of 1.11 ± 0.24 at the very beginning demonstrating the emission of an ensemble of fluorophores for this individual antibody. The *N*_*c*_*/N*_*I,av*_ ratio dropped to 0.77 ± 0.27 after a few hundred milliseconds reflecting the first bleaching step and to 0.49 ± 0.01 after the second bleaching step, which demonstrates the emission of two remaining emitting fluorophores. The *N*_*c*_*/N*_*I,av*_ ratio of 0.19 ± 0.02 measured after the third bleaching step demonstrates the emission of a single fluorophore (**Supplementary Fig. 11b**). Intriguingly, we also observed fluorescence trajectories of higher DOL antibodies that showed clear photon antibunching signs, i.e. *N*_*c*_*/N*_*I,av*_ ratios of 0.2 – 0.3 in aqueous trolox buffer (**Supplementary Figure 14**).

To explain why a few multiple-Al647-labeled antibodies behave partially as single-quantum systems, an efficient nonradiative pathway has to be considered. Depending on the distance between Al647 monomers residing in the on-state (singlet ground and excited state) strong or weak molecular coupling can occur^16,17,49-51,55^. In the event of strong electronic interactions, the Al647 monomers will form non-fluorescent dimers. Consequently, the monomer-dimer equilibrium will control the emitted fluorescence intensity. For weak electronic interactions between the dyes dipole-dipole induced energy transfer can occur^34,35^. Accordingly, the energy transfer efficiency scales with the sixth power of the distance between the dyes. If the distance and orientations of identical dyes are kept constant, the energy transfer efficiency is determined solely be the overlap of the emission and absorption of the dye. Hence, for dyes with small Stokes shift such as carbocyanine dyes the overlap is sufficient to enable so-called homo-transfer or energy hopping, which occurs in some light-harvesting complexes and has also been observed for artificial multichromophoric systems^25-27,57^. In addition, nonradiative energy transfer from one dye residing in the first excited singlet state (S_1_) to another dye residing in the triplet (T_1_) or excited singlet state (S_1_) is possible. If the fluorescence emission of the dye is in resonance with a transition of the second dye into a higher excited triplet (T_n_) or singlet state (S_n_), the excited state of the donor dye is annihilated. Because the process results in only one excited state remaining in the multiple-labeled antibody, it is often referred to as S_1_-T_1_ and S_1_-S_1_ annihilation, respectively^25^. Since the off-state of carbocyanine dyes like Al647 absorb at shorter wavelengths energy transfer from S_1_ to the off-state is unlikely^22,45,46^. Consequently, the excited state energy of multiple-Al647-fluorophores residing simultaneously in the on-state can be annihilated in some cases with optimal interaction distance and orientation of fluorophores. This results in emission characteristics expected for a single emitter.

In a nut shell, multiple-Al647-labeled antibodies behave like multichromophoric systems composed of several dynamically interacting fluorophores that emit at least partially independently in aqueous trolox buffer.

### Photon antibunching of multiple-Al647-labeled antibodies in photoswitching buffer

On the other hand, and in strong contrast all multiple-Al647-labeled antibodies showed clear signatures of single emitters in photoswitching buffer containing 100 mM MEA and oxygen scavenger (**Figs. 4f-i**). The average *N*_*c*_/*N*_*l,av*_ ratios measured for different DOLs of 0.23 ± 0.13 (DOL 1.1), 0.31 ± 0.15 (DOL 2.1), 0.19 ± 0.10 (DOL 4.1), and 0.27 ± 0.11 (DOL 8.3) (**Fig. 4j**) demonstrate that the emission of multiple-Al647-labeled antibodies, i.e. the on-states are dominated by single emitting Al647 fluorophores. These findings answer the first question and explain why multiple-Al647-labeled antibodies enable DOL independent *d*STORM imaging in photoswitching buffer. However, the second question why the DOL does not seem to affect the on- and off-state lifetime still has not been fully and sufficiently answered. That is, the origin of the nearly identical blinking behavior of multiple-Al647-antibodies in photoswitching buffer remains to a large extent puzzling. The data imply that the number of on-events and localizations is independent of the DOL (**Table 1 and Supplementary Table 2**).

Interestingly, by examining the confocal single-molecule data more carefully, we identified fluorescence trajectories of multiple-Al647-labeled antibodies with higher DOL (4.1 and 8.3) that showed a firework of blinking events during the first seconds of irradiation (**Figure 5**). Such frequent blinking of single antibodies cannot be captured in widefield *d*STORM imaging experiments. For *d*STORM imaging the sample has to be irradiated for seconds to minutes before data acquisition can be started to transfer the majority of fluorophores from the on- to the off-state and enable the spatially isolated detection and localization of individual fluorescent probes. In turn this means that higher DOL antibodies exhibit at the very beginning of the experiment shorter off-times and can therefore theoretically more often be localized. However, on the consideration that the first few seconds and blinking events will always be missed, all multiple-Al647-labeled antibodies will behave identical in *d*STORM experiments.

**Figure 5.**
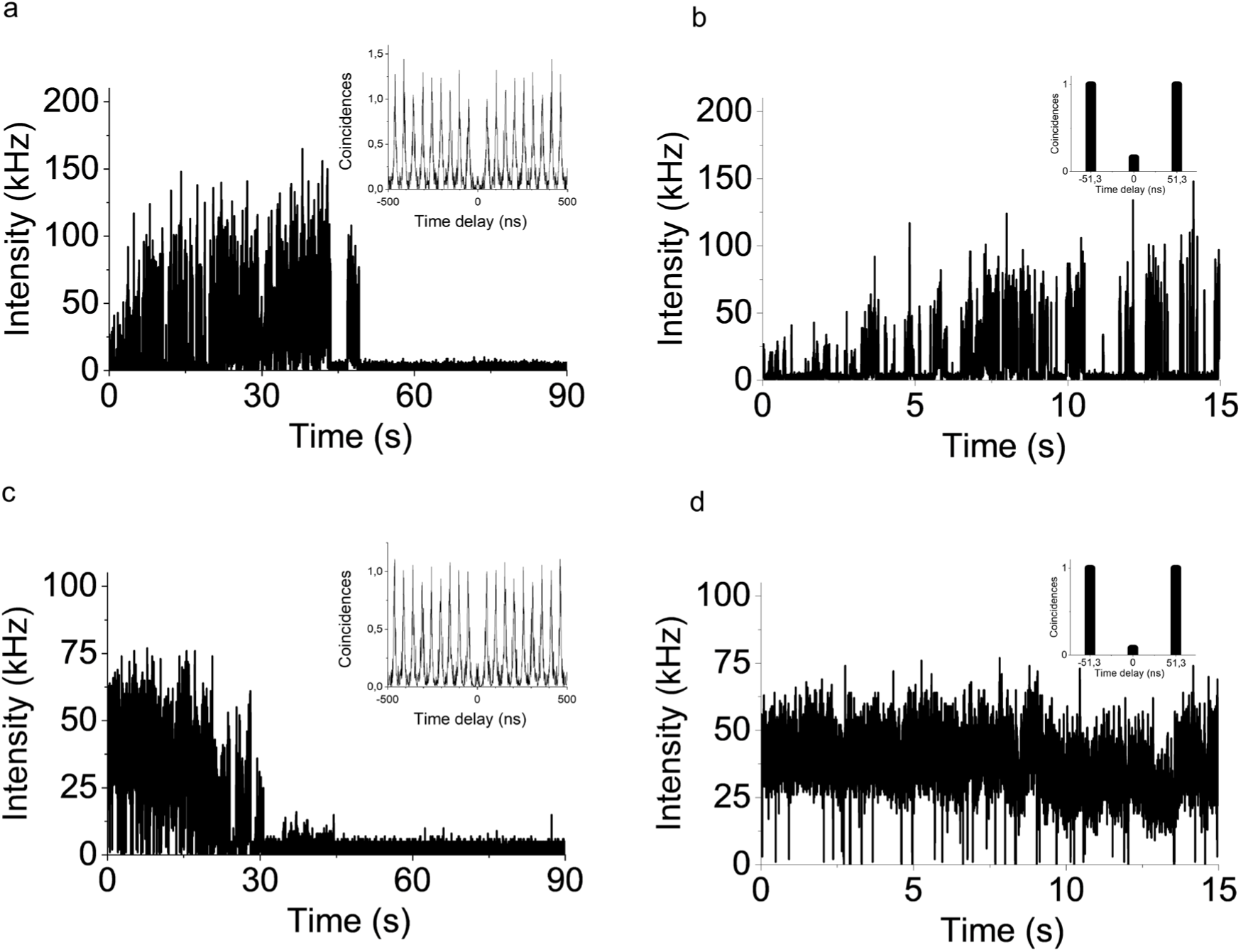
Multiple-Al647-labeled antibodies show frequent blinking at the very beginning of irradiation. (**a**) Fluorescence trajectory of a multiple-Al647-labeled antibody with DOL 4.1 measured in photoswitching buffer (100 mM MEA, oxygen scavenger) excited at 640 nm with 1 kW cm^-2^ (1 ms binning). The inset shows the corresponding normalized interphoton time (coincidence) histogram measured for the entire trajectory. (**b**) Enlarged fluorescence trajectory recorded for the first 15 s highlighting the blinking events. The inset shows the simplified interphoton time (coincidence) histograms containing the zero-time delay and the first lateral peaks. For the entire trajectory and for the first 15 s we determined *N*_*c*_*/N*_*I,av*_ ratios of 0.05 ± 0.01 and 0.16 ± 0.01, respectively, demonstrating emission of a single fluorophore. (**c**) Fluorescence trajectory of a multiple-Al647-labeled antibody with DOL 8.3 measured in photoswitching buffer (100 mM MEA, oxygen scavenger) excited at 640 nm with 1 kW cm^-2^ (1 ms binning). The inset shows the corresponding normalized interphoton time (coincidence) histogram measured for the entire trajectory. (**d**) Enlarged fluorescence trajectory recorded for the first 15 s highlighting the blinking events. The inset shows the simplified interphoton time (coincidence) histograms containing the zero-time delay and the first lateral peaks. For the entire trajectory and for the first 15 s we determined *N*_*c*_*/N*_*I,av*_ ratios of 0.09 ± 0.01 and 0.08 ± 0.01, respectively, demonstrating emission of a single fluorophore.

Our data imply that during the first few seconds of irradiation the equilibrium between the on- and off-state populations of Al647 molecules is adjusted, which results for higher DOL antibodies occasionally in the appearance of fast on/off blinking events during the first seconds. Nevertheless, photon antibunching analysis during the frequent blinking phase at the beginning of the experiments revealed *N*_*c*_*/N*_*I,av*_ ratios of 0.16 ± 0.01 for DOL 4.1 and 0.08 ± 0.01 for DOL 8.3, respectively, demonstrating unequivocally that higher DOL multiple-Al647-labeled antibodies behave like single emitters in photoswitching buffer (**Figs. 5b,d**).

However, in our line of our argumentation there remains an unneglectable possibility that several fluorophores reside simultaneously in the on-state, especially during the first seconds (**Figure 5**). Here again as seen also for some antibodies in trolox buffer (**Supplementary Fig. 14**) the excited state energy of multiple Al647 fluorophores residing simultaneously in the on-state can be annihilated, resulting in emission characteristics expected for a single emitter^25-27,57^.

## Discussion

The DOL of commercially available antibodies has been optimized to 2-5 to minimize quenching effects and retain a high brightness and specific binding of the fluorescent probe^16,19,20^. Normally, single-molecule localization microscopy methods such as *d*STORM use immunostaining for super-resolved visualization of endogenous proteins. However, so far the influence of the DOL of antibodies in *d*STORM experiments has been ignored although the number of fluorophores attached per antibody does not only influence the fluorescence intensity but should also control the blinking behavior of the probe. Our investigations clarify that multiple-Al647-labeled antibodies are very well suited for *d*STORM imaging independent of the DOL. Multiple-Al647-labeled antibodies with DOL 1.1, 2.1, 4.1, and 8.3 exhibit high localization precisions enabling in combination with efficient immunostaining spatial resolutions of ∼ 20 nm (**Table 1, Supplementary Table 2, Figure 3** and **Supplementary Fig. 8**). However, our data also show that higher DOL antibodies tend to bind nonspecifically resulting in higher background signals (**Figure 3** and **Supplementary Figs. 4-7**). In addition, we have shown that the different average fluorescence lifetimes of multiple-Al647-labeled antibodies (**Supplementary Table 1**) can be used advantageously for multi-target FLIM^43^ at the ensemble level (**Figure 2**). Direct comparison of ensemble and single-molecule data recorded from multiple-Al647-labeled antibodies showed that strong electronic interactions between individual fluorophores conjugated to the same antibody are reduced in photoswitching buffer. To explain the observed behavior in aqueous buffer and photoswitching buffer we performed single-molecule fluorescence trajectories and photon antibunching experiments (**Figure 4**). Our data clearly showed that multiple-Al647-antibodies exhibit partially complex emission characteristics indicating weak and strong electronic interactions between fluorophores. This situation completely changes if experiments are performed in photoswitching buffer containing 100 mM MEA and oxygen scavenger. Under these conditions, all multiple-Al647-labeled antibodies behave like single emitters (**Figure 4**), even at the very beginning of irradiation when high DOL antibodies exhibited fast blinking events (**Figure 5**). The observation of non-classical light emitted by multiple-labeled antibodies can be explained by efficient removal of fluorophore monomers in the on-state from the monomer-dimer equilibrium and subsequent accumulation of fluorophores in the long-lived off-state. The shortening of off-state lifetimes measured for high DOL antibodies during the first seconds (**Figure 5**) might be explained by light-induced near-field activation of fluorophores in the off-states. Even if several fluorophores reside simultaneously in the on-state the fluorophore with the lowest excited singlet-state energy (due to small differences in the nanoenvironment) will serve as an ‘emissive trap’ and collect all excited state energy via energy hopping. If this fluorophore eventually photobleaches another fluorophore with the next lowest excited state energy will take over^25-27^. To conclude, multiple-Al647-labeled antibodies exhibit identical blinking performance because the first few seconds with frequent blinking are missed in *d*STORM experiments. Based on our findings the optimal multiple-labeled antibody for *d*STORM, i.e. a kind of ‘super-emitter’, contains several fluorophores separated by 2-4 nm. This inter-fluorophore distance prevents strong electronic interactions of fluorophores such as the formation of non-fluorescent H-dimers but still enables weak electronic interactions via energy transfer. Such a mulichromophoric system would exhibit a higher absorption cross section under circular polarized excitation and due to energy hopping of weakly interacting fluorophores behave like a single emitter in *d*STORM experiments. Such a ‘super-emitter’ could be realized by introducing unnatural amino acids (e.g. TCO-lysine) site-specifically at well-defined positions via genetic code expansion (GCE) into recombinantly expressed antibodies followed by bioorthogonal click labeling with tetrazine-dyes^58,59^.

## Methods

### Antibody labeling

For antibody labeling at different degrees of labeling (DOL) an excess of Alexa Fluor 647-NHS (LifeTech, A20106) and Cy5-NHS (GE-Healthcare, PA15101), respectively, was used. Goat anti-rabbit IgG (Invitrogen, 31212) was used as secondary antibody for the microtubule and single molecule surfaces staining and goat anti-mouse IgG (Sigma-Aldrich, SAB3701063-1) was used for β-tubulin immunostaining in FLIM experiments. Antibody labeling was performed at RT for 4h in labeling buffer (100 mM sodium tetraborat (Fulka, 71999), pH 9.5) following the manufacturers standard protocol. Briefly, 100 µg antibody was reconstituted in labeling buffer using 0.5 ml spin-desalting columns (40K MWCO, ThermoFisher, 87766). Different excesses of Alexa Fluor 647-NHS were used to adjust the DOL. For the goat anti-mouse IgG DOL 0.9 and goat anti-rabbit IgG DOL 1.1 a 2x excess and for goat anti-rabbit IgG DOL 2.1, DOL 4.1, DOL 8.3 a 5x, 20x and 35x excess was used. Additionally, for preparation of single-molecule antibody surfaces, the same rabbit anti-α-tubulin primary antibody (Abcam, ab18251), which was used for the microtubule samples, was labeled with Biotin-NHS (Merck, 203112). Therefore 100 µg of 1g/l primary antibody were treated according to the labeling protocol above using 250 µg of Biotin-NHS. Antibody conjugates were purified and washed up to three times using spin-desalting columns (40K MWCO) in PBS (Sigma-Aldrich, D8537-500ML) to remove excess dyes. Finally, antibody concentration and DOL was determined by UV-vis absorption spectrometry (Jasco V-650).

### Absorption and emission spectra

Steady-state absorption and fluorescence emission spectra were recorded on a V-650 spectrophotometer (Jasco) and a FP-6500 spectrofluorometer (Jasco). Samples were measured in a 0.3 mm path-length fluorescence cuvette (Hellma, 105.251-QS) in PBS (Sigma-Aldrich, D8537-500ML). The temperature was controlled using a Peltier thermocouple set to 25 °C. Concentrations were kept below 1 µM.

### Fluorescence lifetime

The ensemble fluorescence lifetimes (τ) were measured in a 0.3 mm path-length fluorescence cuvette (Hellma, 105.251-QS) on a FluoTime 200 time-resolved spectrometer (PicoQuant, Berlin, Germany) using a pulsed diode laser (635 nm) as the excitation source with a Sepia II module (PicoQuant, Berlin, Germany) and a PicoHarp300 TCSPC module and picosecond event timer (PicoQuant, Berlin, Germany) (80 MHZ, 50 ps pulse length, 8 ps resolution, 10.000 photons in the maximum channel). The results were analyzed with the FluoFit 4.4.0.1 software (PicoQuant, Berlin, Germany). To exclude polarization effects, fluorescence was observed under the magic angle (54.7°). The decay parameters were determined by least-square deconvolution, and their quality was judged by the reduced χ 2 values.

### Cell culture

African green monkey kidney fibroblast-like cells (COS7, Cell Lines Service GmbH, Eppelheim, #605470) were cultured in DMEM (Sigma, #D8062) containing 10 % FCS (Sigma-Aldrich, #F7524), 100 U/ml penicillin and 0.1 mg/ml streptomycin (Sigma-Aldrich, #P4333) at 37 °C and 5 % CO2. Cells were grown in standard T25-culture flasks (Greiner Bio-One).

### Immunostaining protocol

For immunostaining, cells were seeded at a concentration of 2.5 × 10^4^ cells/well into 8 chambered cover glass systems with high performance cover glass (Cellvis, C8-1.5H-N) and stained after 3 hours of incubation at 37 °C and 5 % CO_2_. For microtubule and clathrin immunostaining, cells were washed with pre-warmed (37 °C) PBS (Sigma-Aldrich, D8537-500ML) and permeabilized for 2 min with 0.3 % glutaraldehyde (GA) + 0.25 % Triton X-100 (EMS, 16220 and ThermoFisher, 28314) in pre-warmed (37° C) cytoskeleton buffer (CB), consisting of 10 mM MES ((Sigma-Aldrich, M8250), pH 6.1), 150 mM NaCl (Sigma-Aldrich, 55886), 5 mM EGTA (Sigma-Aldrich, 03777), 5 mM glucose (Sigma-Aldrich, G7021) and 5 mM MgCl_2_ (Sigma-Aldrich, M9272). After permeabilization cells were fixed with a pre-warmed (37° C) solution of 2 % GA for 10 min. After fixation, cells were washed twice with PBS (Sigma-Aldrich, D8537-500ML). After fixation, samples were reduced with 0.1 % sodium borohydride (Sigma-Aldrich, 71320) in PBS for 7 min. Cells were washed three times with PBS (Sigma-Aldrich, D8537-500ML) before blocking with 5 % BSA (Roth, #3737.3) for 30 min. Subsequently, microtubule samples were incubated with 2 ng/µl rabbit anti-α-tubulin primary antibody (Abcam, #ab18251), the clathrin samples were incubated with 2 ng/µl rabbit anti-clathrin primary antibody (Abcam, #ab21679) and mouse anti-β-tubulin primary antibody (Sigma-Aldrich, T8328) in blocking buffer for 1 hour. After primary antibody incubation, cells were rinsed with PBS (Sigma-Aldrich, D8537-500ML) and washed twice with 0.1 % Tween20 (ThermoFisher, 28320) in PBS (Sigma-Aldrich, D8537-500ML) for 5 min. After washing, cells were incubated in blocking buffer with 4 ng/µl of multiple-labeled goat anti-rabbit IgG secondary antibodies (Invitrogen, 31212) with different DOLs for the α-tubulin, 4 ng/µl of the multiple-labeled goat anti-mouse IgG secondary antibodies (Sigma-Aldrich, SAB3701063-1) for the β-tubulin samples and 4 ng/µl of the multiple-labeled goat anti-rabbit IgG secondary antibody DOL 8.3 (Invitrogen, 31212) for the clathrin samples for 45 min. After secondary antibody incubation, cells were rinsed with PBS (Sigma-Aldrich, D8537-500ML) and washed twice with 0.1 % Tween20 (ThermoFisher, 28320) in PBS (Sigma-Aldrich, D8537-500ML) for 5 min. After washing, cells ware fixed with 4% formaldehyde (Sigma-Aldrich, F8775) for 10 min and washed three times in PBS (Sigma-Aldrich, D8537-500ML) prior to imaging.

### Single molecule surface preparation

For preparation of double-stranded oligonucleotide surfaces, custom-designed single stranded oligonucleotides (IBA, HPLC-grade) containing a 3’-biotinylated sequence (5’-3’: GGGGTATGTGTGTGGGG[BtnTg]) and a 3’-Al647 labeled sequence (5’-3’: CCCCACACACATACCCC[Al647]) were used. For oligonucleotide hybridization, 50 µl of a 10^− 6^ M biotinylated and 10^−7^ M dye-coupled oligonucleotide solution were co-incubated to anneal the oligonucleotides at 80 °C using a Thermocycler (BioRad, C1000 ThermoCycler), followed by sample cooling to 4 °C with -1 °C/min. For the preparation of oligonucleotide single-molecule surfaces, 8 chambered cover glass systems with high performance cover glass (Cellvis, C8-1.5H-N) were used. The glass surfaces were washed once with PBS (Sigma-Aldrich, D8537-500ML) before treatment with 2 % Hellmanex (Hellma, 9-307-011-4-507) for 1 hour. After washing the chambers three times with PBS (Sigma-Aldrich, D8537-500ML), the surfaces were incubated with 1 M KOH (Fulka, 06005) for 20 min. After alkaline treatment, the chambers were washed with PBS (Sigma-Aldrich, D8537-500ML). Afterwards, the surfaces were incubated with 10 % polyethylenglycol 400 (Fulka, 81170) for 4 hours at RT. The surfaces were rinsed with PBS (Sigma-Aldrich, D8537-500ML) before incubating the chambers with blocking solution (10 g/l BSA (ThermoFisher, 30036578) + 0.01 g/l BSA-Biotin (ThermoFisher, 29130) in PBS) overnight at 4 °C. In the following, the chambers were washed three times with PBS (Sigma-Aldrich, D8537-500ML) before incubation with 0.1 g/l Neutravidin (ThermoFisher, 31050) in in PBS (Sigma-Aldrich, D8537-500ML) for 20 min. The surfaces were washed three times with PBS (Sigma-Aldrich, D8537-500ML) and incubated with 10^−10^ M of the annealed oligonucleotides in PBS (Sigma-Aldrich, D8537-500ML) for 1 min. For antibody surfaces, the chambers were treated analogous to the double-stranded oligonucleotide surfaces up to the addition of blocking solution with BSA/BSA-Biotin + Neutravidin. After washing three times with PBS (Sigma-Aldrich, D8537-500ML), the surfaces were incubated with 1 ng/ µl biotinylated alpha-tubulin (Abcam, ab18251) in PBS for 1 min. After washing with PBS (Sigma-Aldrich, D8537-500ML) the chambers were incubated with 0.5 ng/ µl of the custom-labeled goat anti-rabbit IgG secondary antibodies (Invitrogen, 31212) with different degrees of labeling in PBS for 1 min. The prepared samples were washed at least three times in PBS (Sigma-Aldrich, D8537-500ML) prior to imaging.

### *d*STORM imaging

Super-resolution imaging was performed using an inverted wide-field fluorescence microscope (IX-71; Olympus). For excitation of Alexa Fluor 647, a 641 nm diode laser (Cube 640-100C, Coherent), in combination with a clean-up filter (Laser Clean-up filter 640/10, Chroma) was used. The laser beam was focused onto the back focal plane of the oil-immersion objective (60×, NA 1.45; Olympus). Emission light was separated from the illumination light using a dichroic mirror (HC 560/659; Semrock) and spectrally filtered by a bandpass filter (HC697/75 and LP647; Semrock). Images were recorded with an electron-multiplying CCD camera chip (iXon DU-897; Andor). Pixel size for data analysis was measured to 134 nm. For each *d*STORM measurement, 75,000 images with an exposure time of 8 ms (frame rate 125 Hz) and irradiation intensity of 2 kW cm^-2^ were recorded. Microtubules and single-molecule surfaces were imaged by highly inclined and laminated optical sheet (HILO)-illumination and TRIF microscopy, respectively. Experiments were performed in PBS-based photoswitching buffer containing 100 mM β-mercaptoethylamine (MEA, Sigma-Aldrich) and an oxygen scavenger system (5 % (w/v) glucose, 10 U/ml glucose oxidase and 200 U/ml catalase) adjusted to pH 7.6. For dual excitation measurements with 640 nm and UV irradiation, a 405 nm diode laser (Cube 405-100C, Coherent), in combination with a clean-up filter (Laser Clean-up filter 405/20, Chroma) was used. In addition, for excitation with circular polarized light, a quarter-wave plate (Thorlabs, SAQWP05M) was mounted. All *d*STORM results were analyzed with rapidSTORM3.3. The localization precision was calculated according to Mortensen *et al*.^45^.

### Fluorescence Correlation Spectroscopy (FCS)

FCS measurements were performed on a custom-built confocal inverse microscope (Zeiss Axiovert 100 TV) equipped with a 640 nm diode laser (Coherent Cube). The laser beam was coupled into an oil-immersion objective (Zeiss Plan Apochromat, 63×, NA 1.4) via a dichroic beam splitter (Omega Optics 645DLRP). The antibodies of different DOL were diluted to 1 nM in PBS and transferred onto a microscope slide and covered by a coverslip. Sample temperature was set to 25 °C. For each sample, at least three individual ACFs with 5 min measurement time were recorded.

### Fluorescence Lifetime Imaging Microscopy (FLIM)

All fluorescence lifetime images, single-molecule trajectories and photon antibunching measurements were performed on a MicroTime200 (PicoQuant, Berlin, Germany) time-resolved confocal fluorescence microscope setup consisting of a FLIMbee galvo scanner (PicoQuant, Berlin, Germany), an Olympus IX83 microscope including an oil-immersion objective (60×, NA 1.45; Olympus), 2 single photon avalanche photodiodes (SPAD) (Excelitas Technologies, 75154 K3, 75154 L6) and a TimeHarp300 dual channel board. Fro pulsed excitation a white-light laser (NKT photonics, superK extreme) was coupled into the MicroTime200 system via a glass fiber (NKT photonics, SuperK FD PM, A502-010-110). A 100 µm pinhole was used for all measurements. The emission light was split onto the SPADs using a 50:50 beamsplitter (PicoQuant, Berlin, Germany). To filter out after glow effects of the SPADs used as well as scattered and reflected light, 2 identical bandpass filters (ET700/75 M, Semrock, 294808) were installed in front of the SPADs. The measurements were performed and analyzed with the SymPhoTime64 software (PicoQuant, Berlin, Germany). The microtubule and clathrin measurements were performed with an irradiation intensity of ∼0.5 kW cm^-2^ in T3 mode with 25 ps time-resolution, whereas all photon antibunching measurements and single-molecule trajectories were performed with ∼1 kW cm^-2^ in T2 mode. For photon antibunching experiments, the Sync cable was disconnected and replaced by the SPAD 2 cable.

### Photon antibunching measurements

The data in the interphoton time histograms can be quantified for the purpose of determining the number of independent emitters by determining the ratio of the number of photons in the central peak, *N*_*c*_, to the average number in the neighboring lateral peaks, *N*_*l,av*_. Ensemble antibunching measurements show that the number of photon pairs detected in the neighboring peaks decreases at large interphoton times but is nearly constant for very short times, i.e. in the first neighboring peaks. For determination of *N*_*l,av*_, we used the average number of events in the nearest 8 peaks, 4 to each side of the zero-time peak.

## Data availability

All data that support the findings described in this study are available within the manuscript and the related supplementary information. Additional information is available from the corresponding author upon reasonable request.

## Acknowledgement

We thank L. Behringer-Pliess, E. Maier and I. Simeonov for cell culture preparation. This work was supported by the German Research Foundation (DFG, SA829/19-1) and the European Regional Development Fund (EFRE project “Center for Personalized Molecular Immunotherapy”).

## Author contributions

D.H. and M.S. conceived and designed the project. M.S. supervised the project. D.H. performed all labeling and imaging experiments. D.H. and G.B. performed data analysis. D.H., G.B. and M.S. wrote the manuscript.

## Additional information

Supplementary information is available for this paper at:

Reprints and permissions information is available at www.nature.com/reprints.

Correspondence and requests for materials should be addressed to M.S.

## Competing financial interests

The authors declare no competing financial interests.

## Supporting Information

**Supplementary Table 1.** Spectroscopic characteristics of multiple-Al647-labeled antibodies measured in PBS, pH 7.4. Excitation was performed at 640 nm, the emission was recorded at the emission maximum. The decay parameters were determined by least-squares deconvolution, and their quality was judged by the reduced *χ*^2^ values and the randomness of the weighted residuals (*χ*^2^ = 0.9 – 1.1). In the case that a monoexponential model was not adequate to describe the measured decay, a multiex-ponential model was used to fit the decay *(τ*_av_ = *τ*_1_a_1_ + *τ*_2_a_2_). For free Al647 we measured a monoexponential fluorescence decay with a lifetime of 1.09 ns.

| DOL | $\lambda_{abs,max}$ (nm) | $\lambda_{em,max}$ (nm) | $\tau_1$ (ns)/$a_1$ | $\tau_2$ (ns)/$a_2$ | $\tau_{av}$ (ns) |
| --- | --- | --- | --- | --- | --- |
| 1.1 | 651 | 667 | 1.65 / 0.65 | 0.96 / 0.35 | 1.29 |
| 2.1 | 651 | 668 | 1.53 / 0.76 | 0.64 / 0.24 | 1.28 |
| 4.1 | 651 | 670 | 1.33 / 0.54 | 0.29 / 0.46 | 0.74 |
| 8.3 | 651 | 672 | 1.16 / 0.28 | 0.20 / 0.72 | 0.38 |

**Supplementary Table 2.**
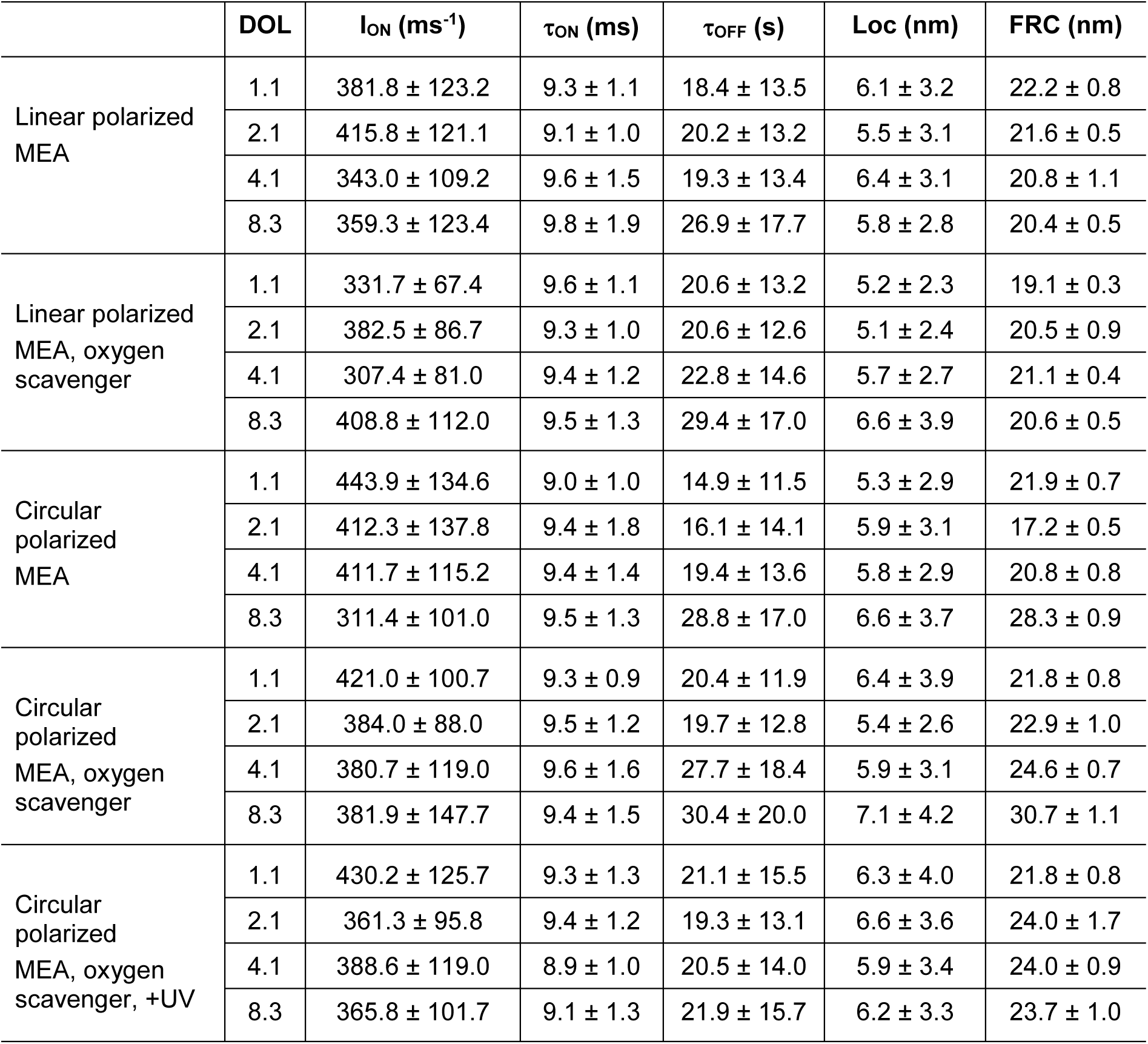
On-state fluorescence intensity in photons ms^-1^ (I_ON_), on-state lifetime (*τ*_ON_) in milliseconds, off-state lifetime (*τ*_OFF_) in seconds, average *xy* localization precision in nanometers (Loc)^45^, and spatial resolution (FRC)^46^ determined from *d*STORM data of COS7 microtubules immunostained with multiple-Al647-labeled antibodies. Data were recorded under different irradiation and buffer compositions and conditions: linear or circular polarized in 100 mM MEA, PBS, pH 7.6 in the absence and presence of oxygen scavenger. Samples were excited at 641 with an irradiation intensity of 2 kW cm^-2^ without or with additional 405 nm with 8 W cm^-2^ (+UV). Detection was performed at a frame rate of 125 Hz. Subsequent blink events of the same antibody were histogrammed and the average parameters determined. The average parameters of each antibody were then histogrammed to determine the mean values ± standard deviation (s.d.) (**Supplementary Figs. 4-7**).

**Supplementary Figure 1.**
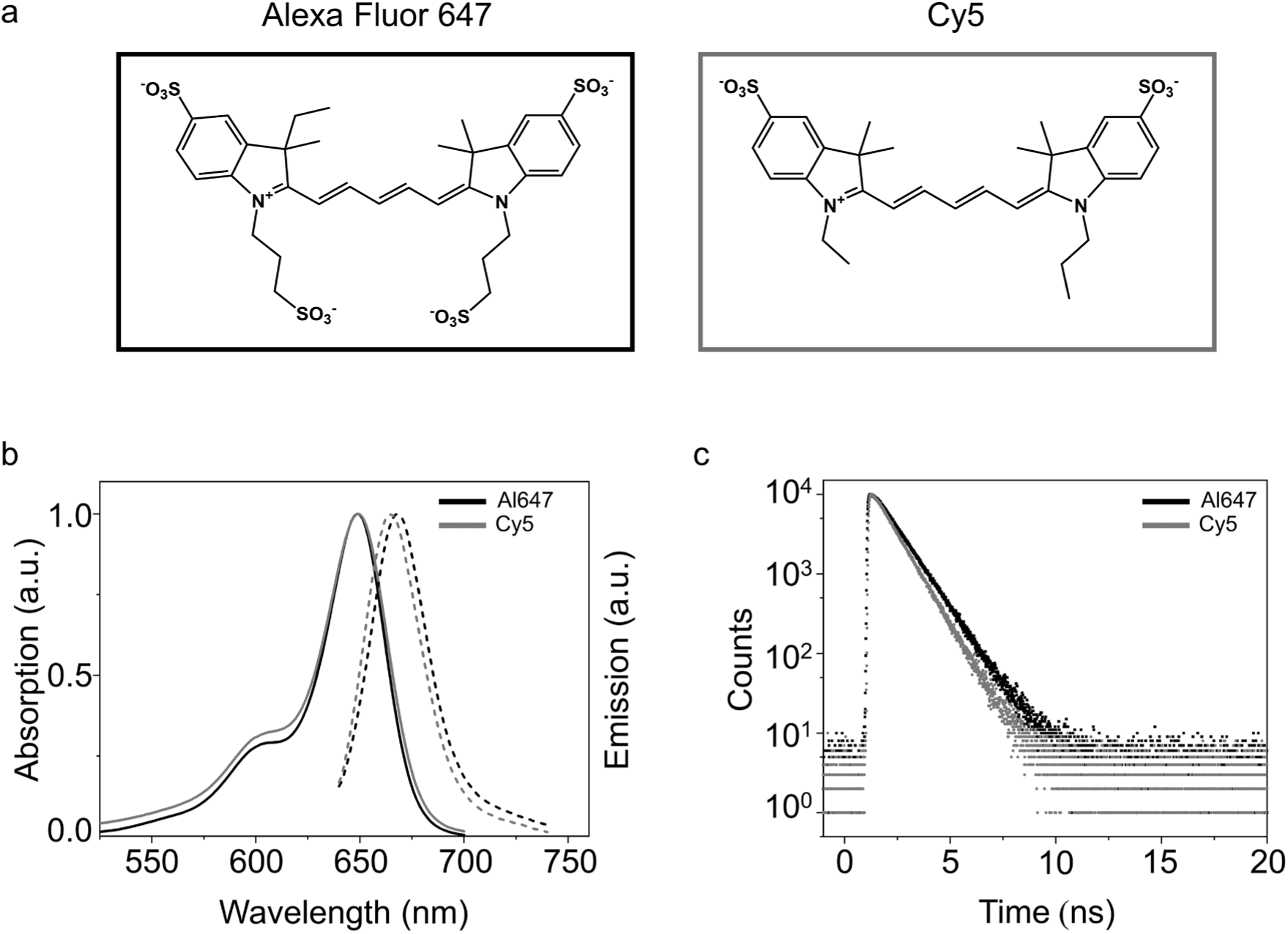
**(a)** Molecular structures of Alexa Fluor 647 and Cy5. (b) Normalized absorption (solid lines) and emission (dashed lines) spectra of Alexa Fluor 647 and Cy5 in PBS, pH 7.4. (**c**) Monoexponential fluorescence decays of Alexa Fluor 647 and Cy5 in PBS, pH 7.4 with lifetimes of 1.09 ns and 0.99 ns.

**Supplementary Figure 2.**
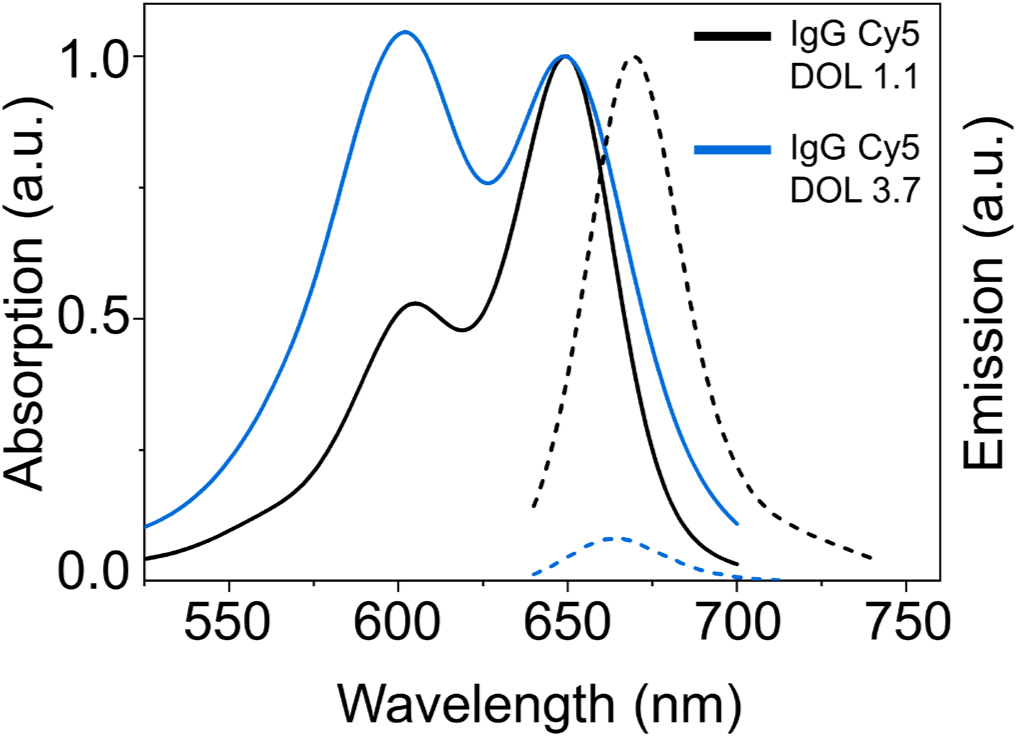
Normalized absorption (solid lined) and relative fluorescence emission (dashed lines) spectra of multiple-Cy5-labeled antibodies with DOL 1.1 (black) and 3.7 (blue).

**Supplementary Figure 3.**
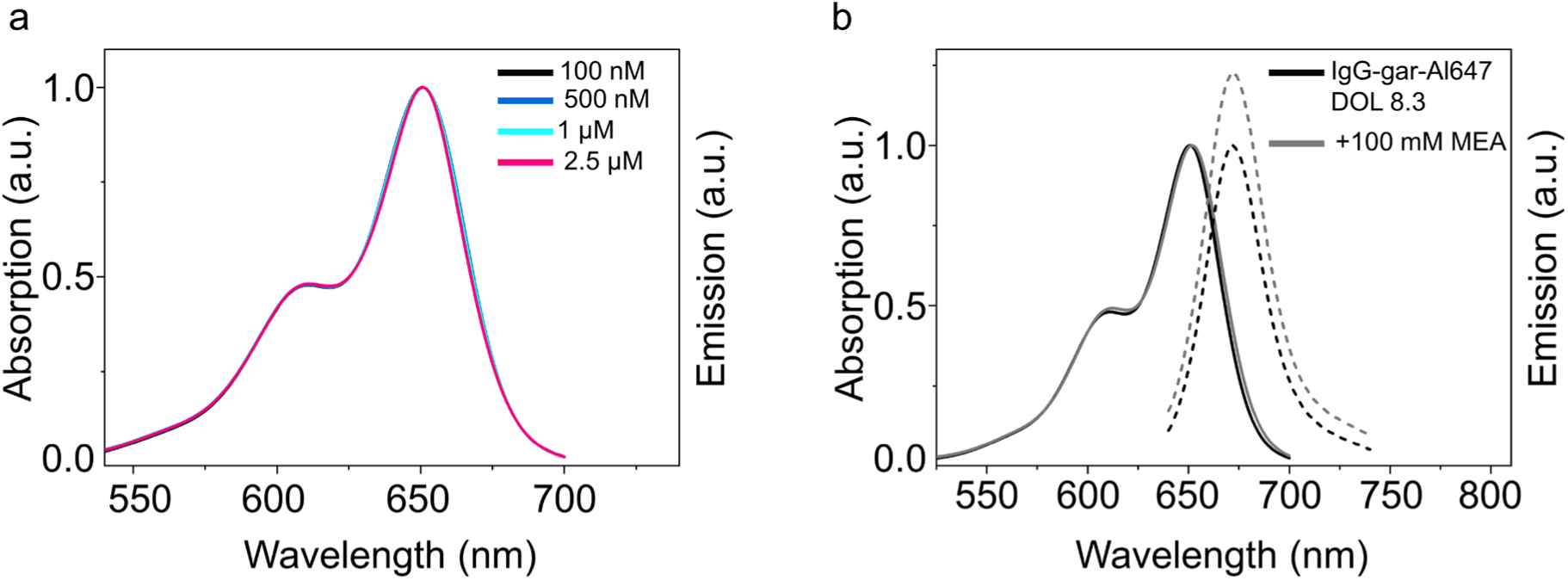
(**a**) Normalized absorption spectra of multiple-Al647-labeled secondary antibodies (DOL 8.3) measured at different concentrations in PBS, pH 7.4. (**b**) Normalized absorption (solid lines) and relative fluorescence emission spectra (dashed lines) of multiple-Al647-labeled secondary antibodies (DOL 8.3) in photoswitching buffer containing 100 mM MEA.

**Supplementary Figure 4.**
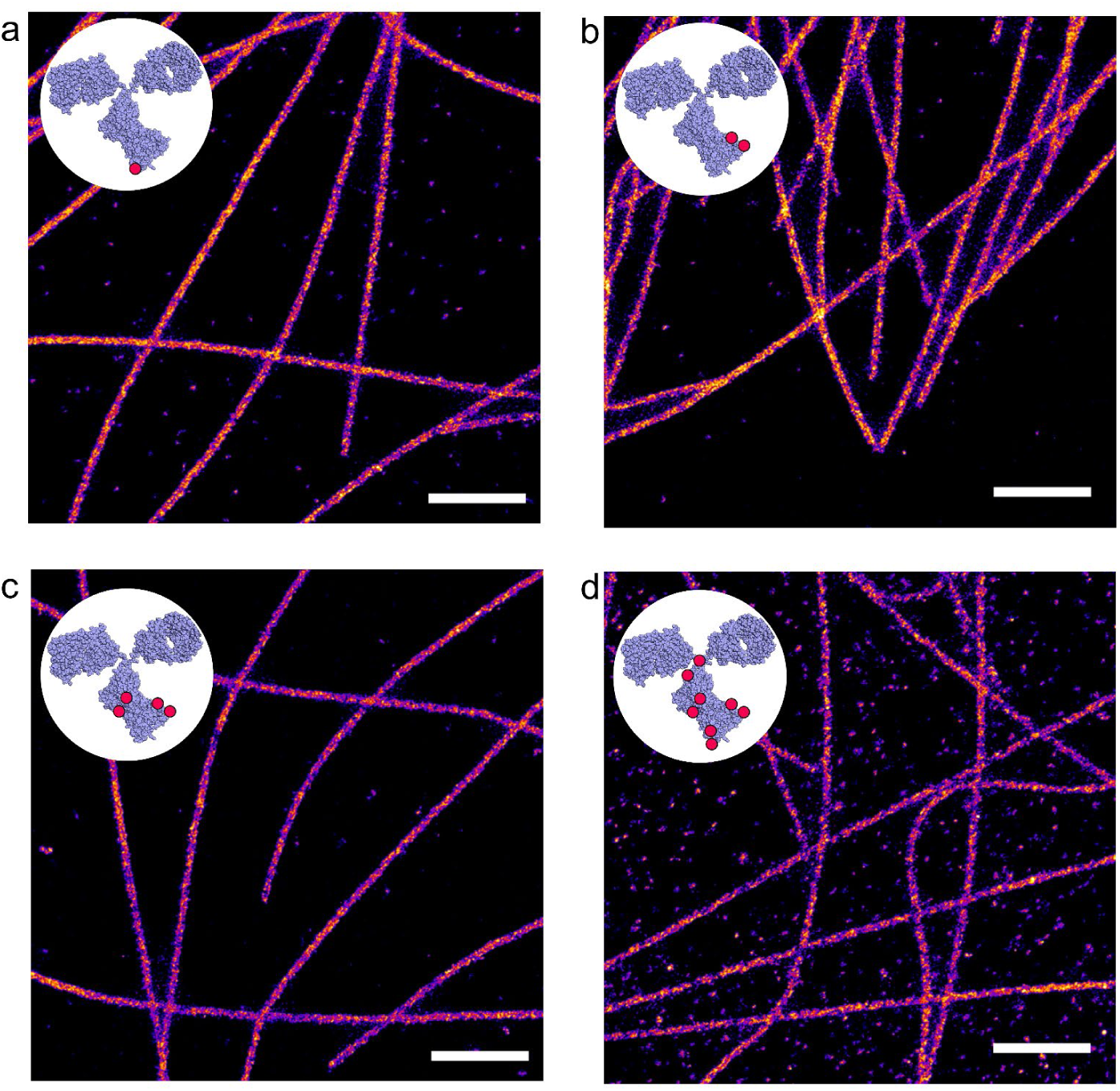
*d*STORM images of microtubules in COS7 cells labeled with secondary multiple-Al647-labeled antibodies with (**a**) DOL 1.1, (**b**) DOL 2.1, (**c**) DOL 4.1 and (**d**) DOL 8.3 recorded under linear polarized laser excitation at 641 nm with an irradiation intensity of 2 kW cm^-2^. Buffer conditions: 100 mM MEA without oxygen scavenging, pH 7.6 Scale bars, 1 µm.

**Supplementary Figure 5.**
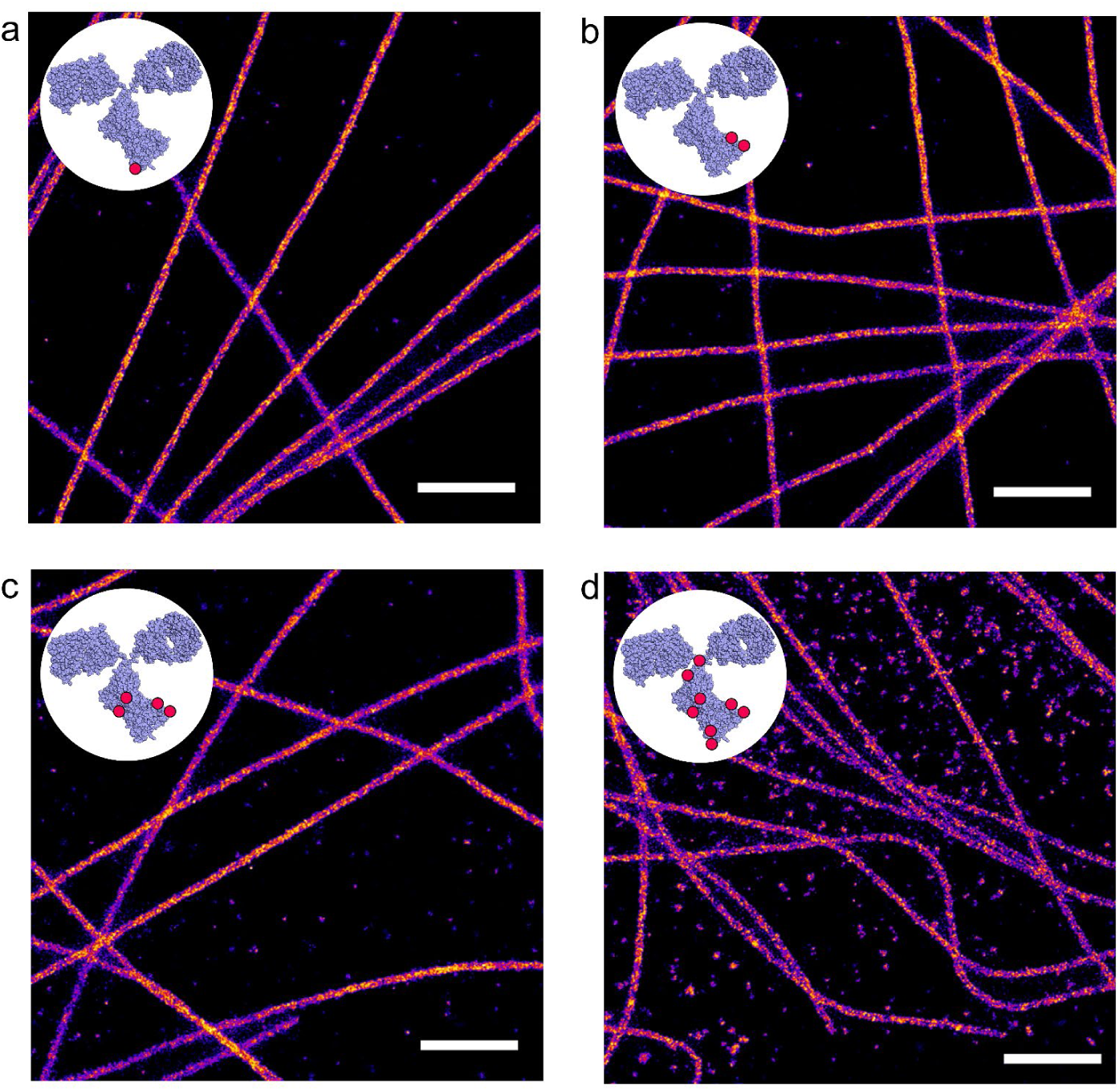
*d*STORM images of microtubules in COS7 cells labeled with secondary multiple-Al647-labeled antibodies with (**a**) DOL 1.1, (**b**) DOL 2.1, (**c**) DOL 4.1 and (**d**) DOL 8.3 recorded under linear polarized laser excitation at 641 nm with an irradiation intensity 2 kW cm^-2^. Buffer conditions: 100 mM MEA with oxygen scavenging, pH 7.6. Scale bars, 1 µm.

**Supplementary Figure 6.**
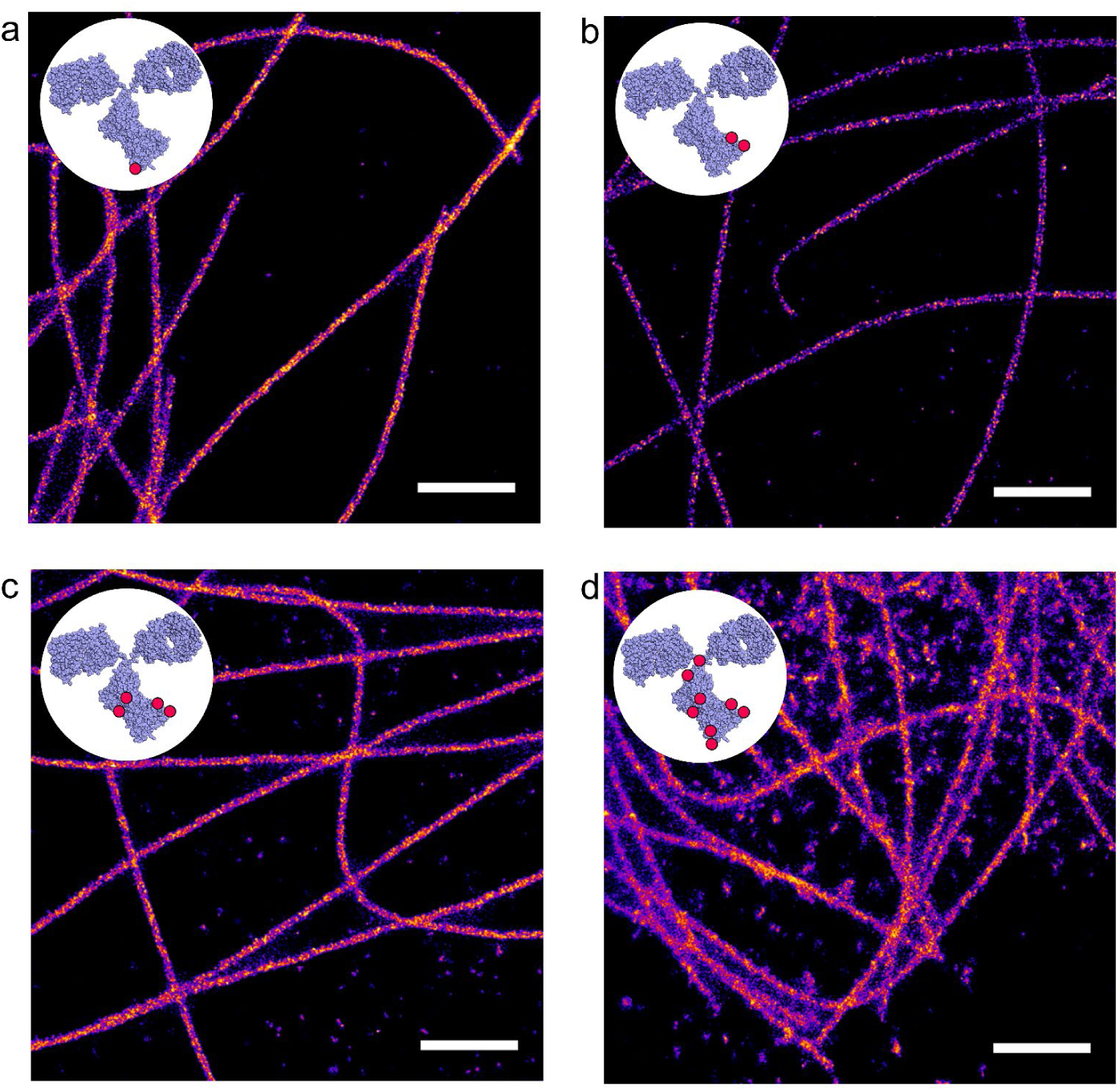
*d*STORM images of microtubules in COS7 cells labeled with secondary multiple-Al647-labeled antibodies with (**a**) DOL 1.1, (**b**) DOL 2.1, (**c**) DOL 4.1 and (**d**) DOL 8.3 recorded under circular polarized laser excitation at 641 nm with an irradiation intensity 2 kW cm^-2^. Buffer conditions: 100 mM MEA without oxygen scavenging, pH 7.6. Scale bars, 1 µm.

**Supplementary Figure 7.**
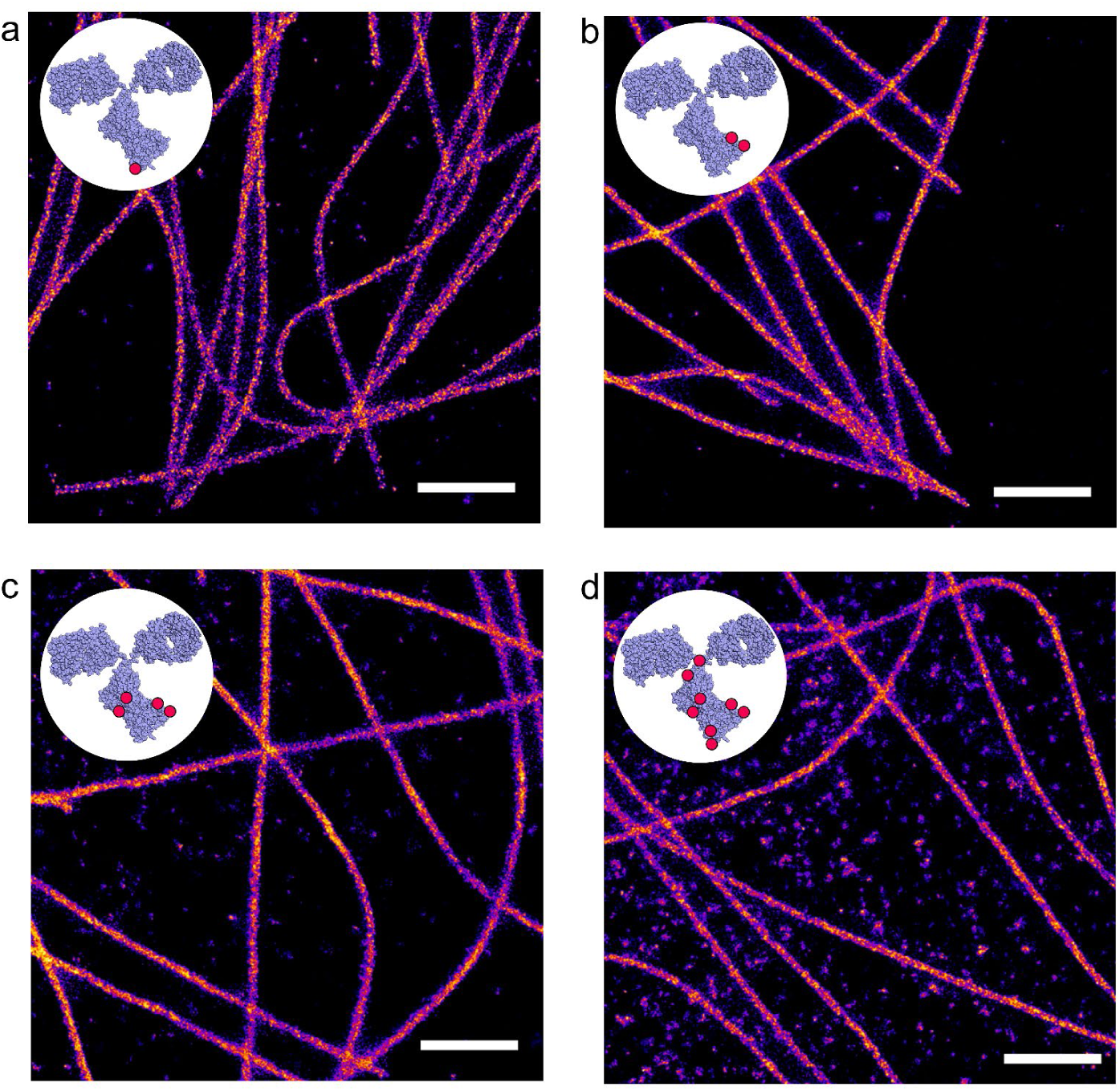
*d*STORM images of microtubules in COS7 cells labeled with secondary multiple-Al647-labeled antibodies with (**a**) DOL 1.1, (**b**) DOL 2.1, (**c**) DOL 4.1 and (**d**) DOL 8.3 recorded under circular polarized laser excitation at 641 nm with an irradiation intensity of 2 kW cm^-2^ and 405 nm with 8 W cm^-2^. Buffer conditions: 100 mM MEA with oxygen scavenging, pH 7.6. Scale bars, 1 µm.

**Supplementary Figure 8.**
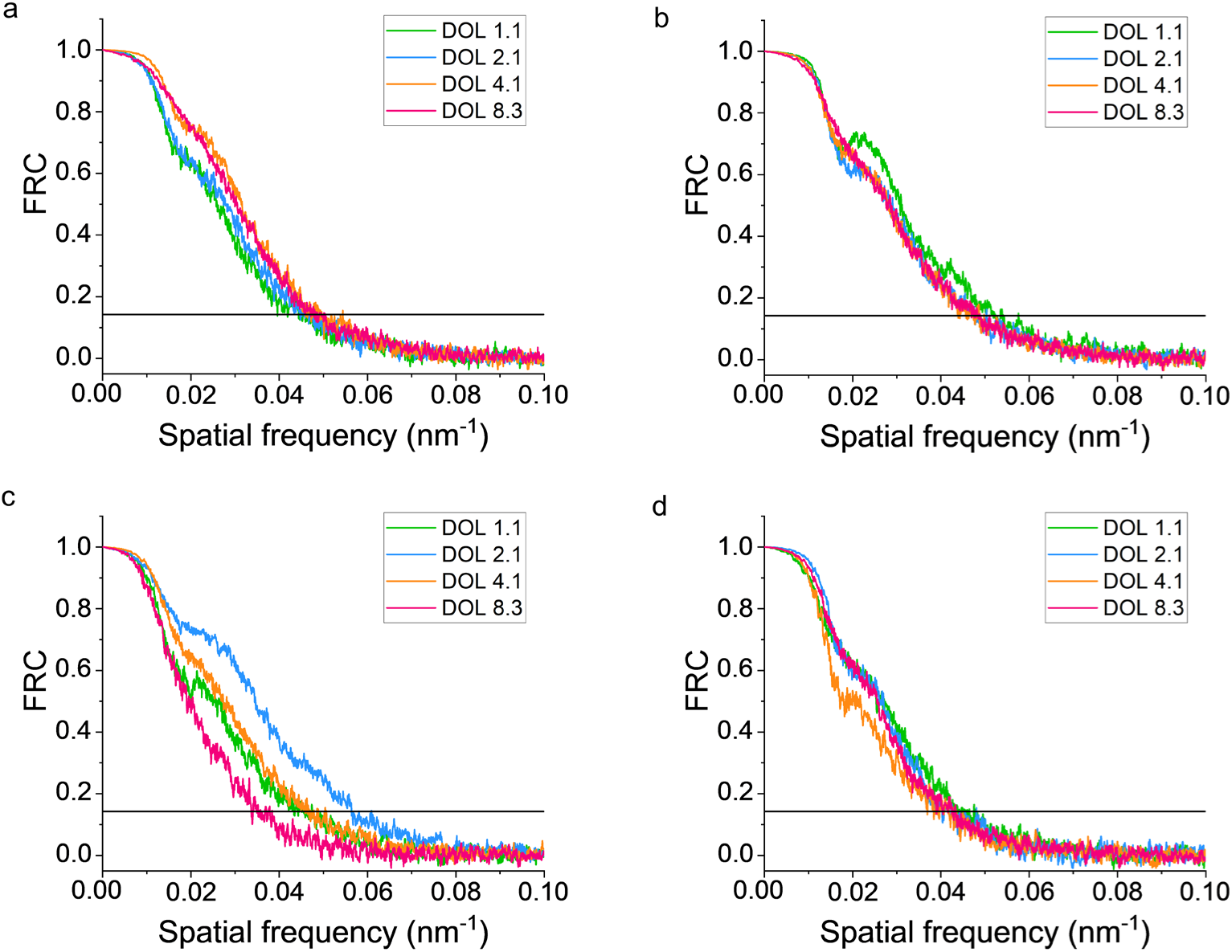
Fourier ring correlation (FRC) resolution estimation of *d*STORM microtubule images of COS7 cells immunostained with multiple-Al647-labeled secondary antibodies with different DOL imaged under different experimental conditions: (**a**) Linear polarized excitation at 641 nm (2 kW cm^-2^) in 100 mM MEA without oxygen scavenger, pH 7.6 (**Supplementary Fig. 4**), (**b**) linear polarized excitation at 641 nm (2 kW cm^-2^) in 100 mM MEA with oxygen scavenger, pH 7.6 (**Supplementary Fig. 5**), (**c**) circular polarized excitation at 641 nm (2 kW cm^-2^) in 100 mM MEA without oxygen scavenger, pH 7.6 (**Supplementary Fig. 6**), (**d**) circular polarized excitation at 641 nm (2 kW cm^-2^) and 405 nm (8 W cm^-2^) in 100 mM MEA with oxygen scavenger, pH 7.6 (**Supplementary Fig. 7**). The calculated values are summarized in **Supplementary Table 2**.

**Supplementary Figure 9.**
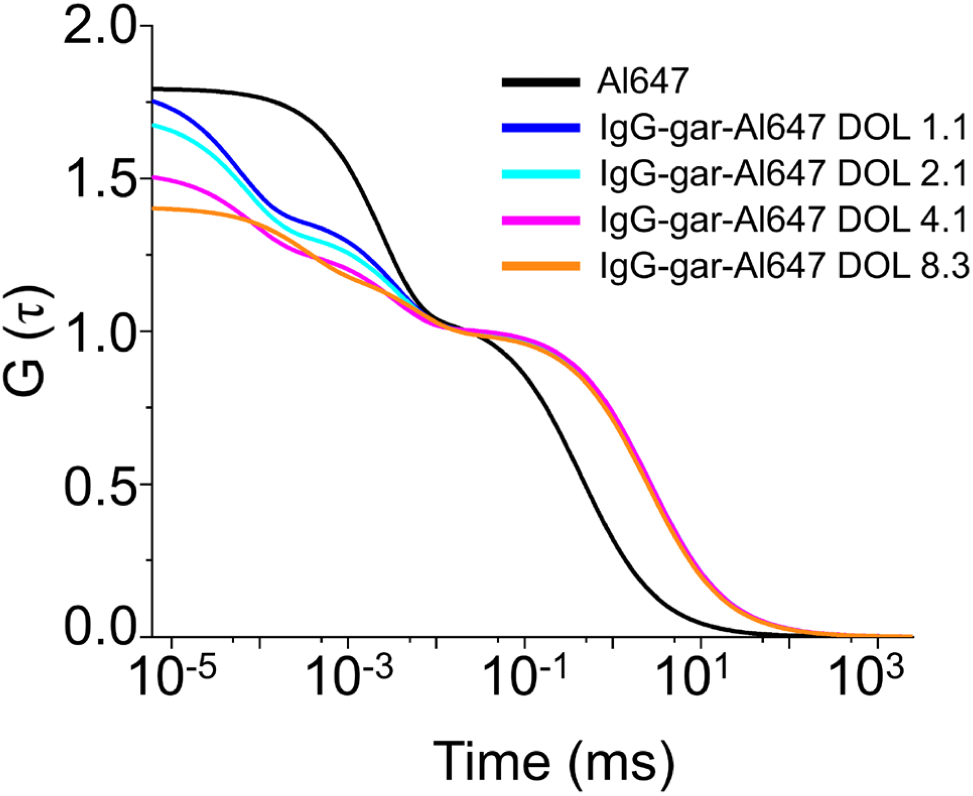
FCS-curves of free Al647 and multiple-Al647-labeled antibodies measured in PBS, pH 7.4.

**Supplementary Figure 10.**
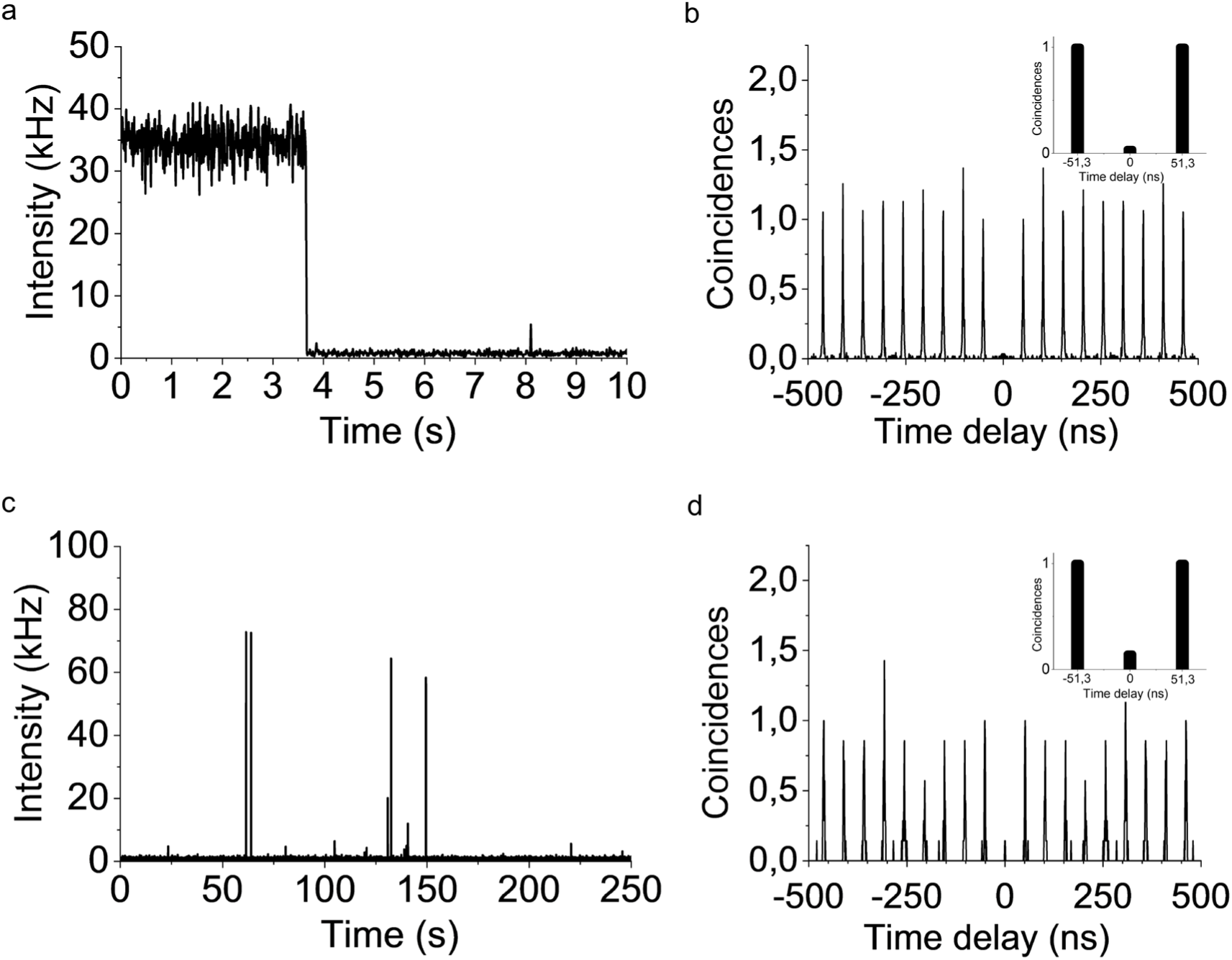
Fluorescence trajectory and coincidence data for a Cy5-DNA construct with one Cy5 molecule measured in trolox buffer. (**a**) Fluorescence intensity as a function of time and (**b**) normalized interphoton distance (coincidence) histogram measured in PBS, pH 7.4. Excitation was performed at 640 nm with a repetition rate of 19.5 MHz. The inset shows only the two first lateral peaks in the time range around the zero peak, which corresponds to photon pairs that were induced by the same laser pulse. (**c**) Fluorescence intensity as a function of time and (**d**) normalized interphoton distance (coincidence) histogram measured in photoswitching buffer containing 100 mM MEA and oxygen scavenger, pH 7.6. For clarity only the central and two first lateral coincidence peaks are shown in the inset. For the determination of *N*_*l,av*_ we used the average number of coincidence events detected in the nearest 8 peaks, 4 to each side of the zero-time peak. The *N*_*c*_/*N*_*l,av*_ ratio for these data is 0.03 ± 0.004 and 0.18 ± 0.04 in trolox and photoswitching buffer, respectively, demonstrating the emission of single emitters. For single molecules the central peak vanishes almost completely since only one photon can be emitted by the molecule after successful excitation.

**Supplementary Figure 11.**
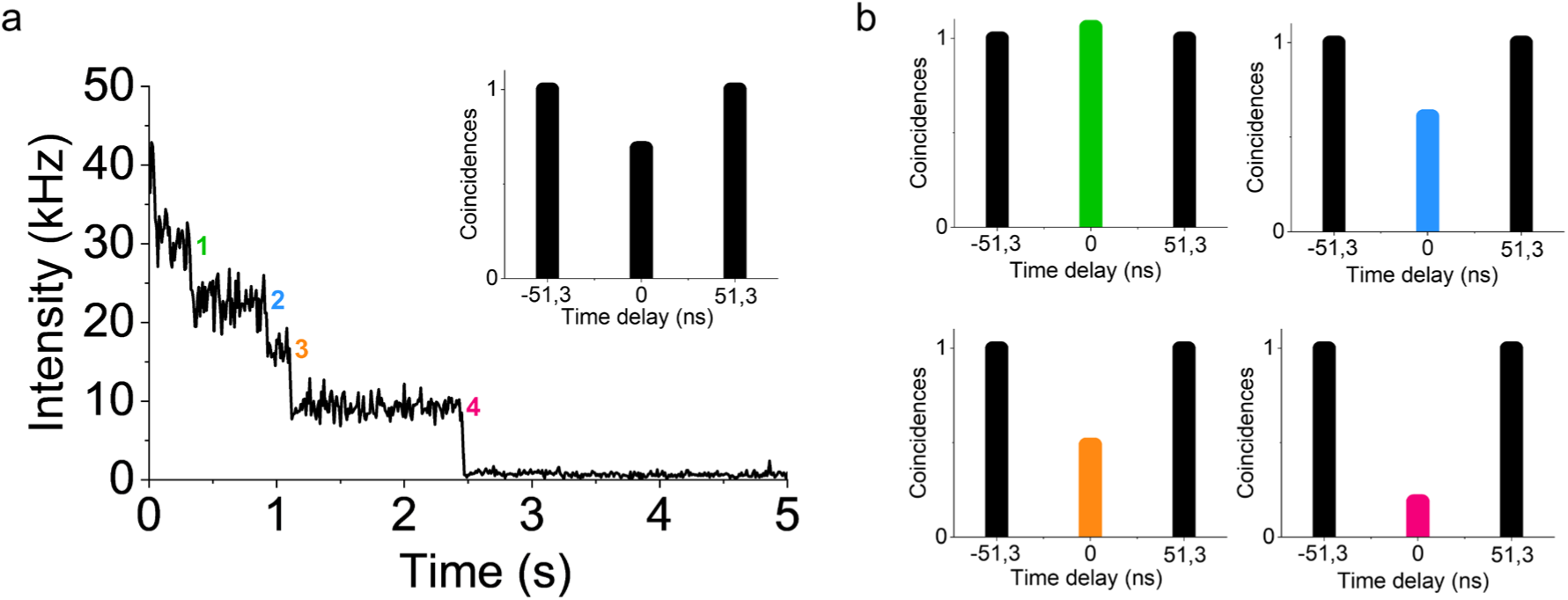
(**a**) Single-molecule fluorescence trajectory of a multi-AL647-labeled antibody, DOL 4.1 in trolox buffer. The inset shows the normalized *N*_*c*_*/N*_*I,av*_ ratio of the antibunching signal measured for the entire fluorescence intensity trajectory. (**b**) Normalized *N*_*c*_*/N*_*I,av*_ ratios recorded for the four different photobleaching steps. The central coincidence peak at zero is color-coded according to the individual bleaching steps shown in (a). The coincidence histograms exhibit a *N*_*c*_*/N*_*I,av*_ ratio of 1.11 ± 0.24 at the beginning of the trajectory demonstrating emission of an ensemble of fluorophores and drops then stepwise down to 0.19 ± 0.02 after the third bleaching step indicating the presence of a single emitter.

**Supplementary Figure 12.**
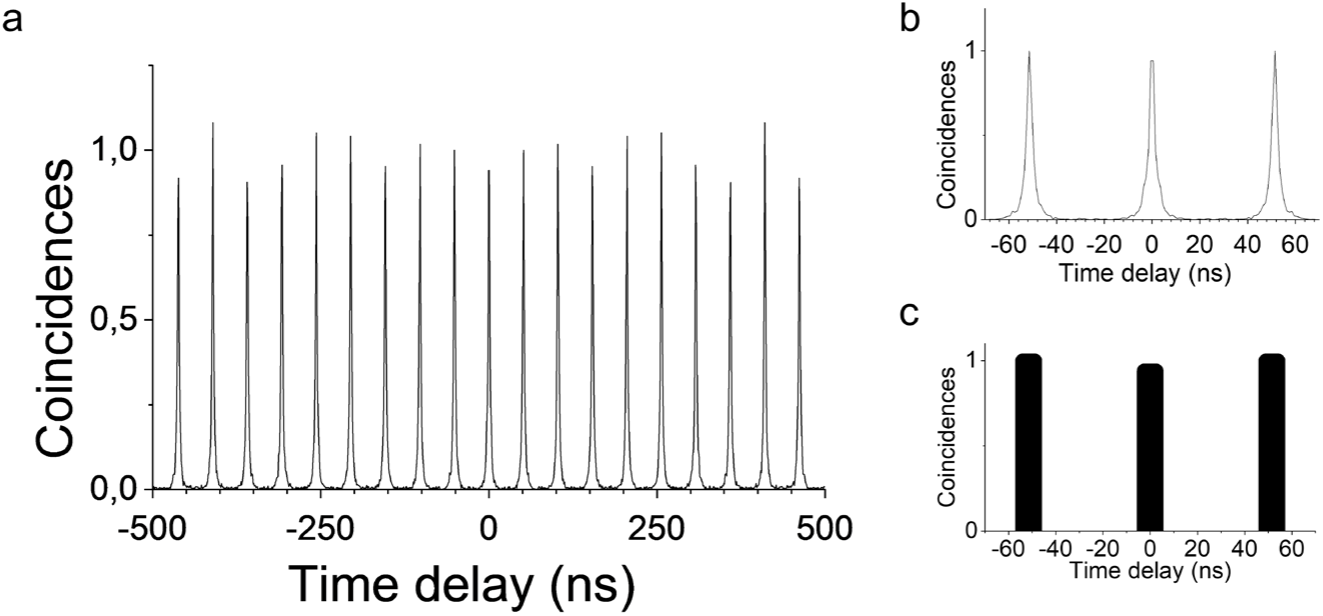
(**a**) Normalized interphoton distance (coincidence) histogram measured from a 200 nm fluorescent microsphere (fluorosphere) parked in the laser focus in PBS, pH 7.4. Excitation was performed at 640 nm with a repetition rate of 19.5 MHz. Hence coincidences events are separated by 51.3 ns. As expected for a Poissonian light source, the obtained peak pattern clearly demonstrates the classical behavior of the radiation field emitted by an ensemble of molecules, i.e. the central peak is identical in intensity to the lateral ones. (**b**) Coincidence histogram showing only the central and the first lateral peaks. (**c**) Simplified presentation of the photon pairs detected in the central, zero-time delay and first lateral peaks at a delay time of 51.3 ns.

**Supplementary Figure 13.**
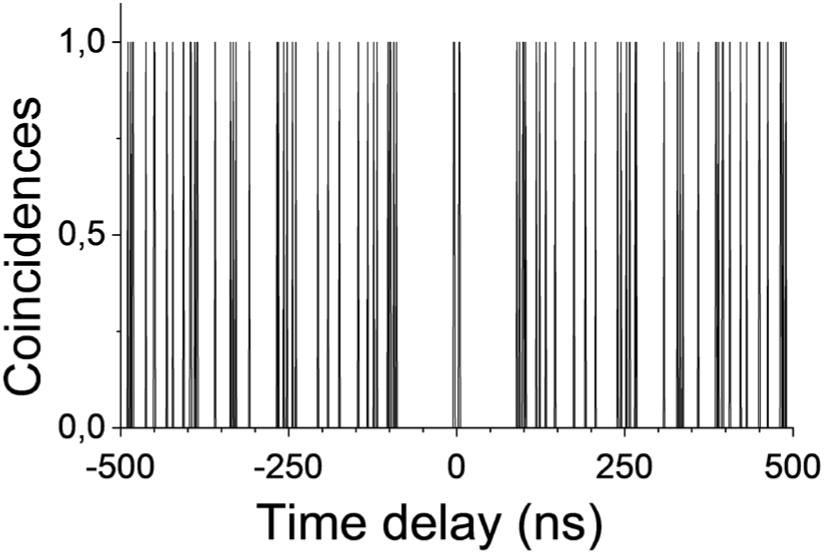
Normalized interphoton distance (coincidence) histogram recorded from the background signal, showing accidental coincidences due to background-background events.

**Supplementary Figure 14.**
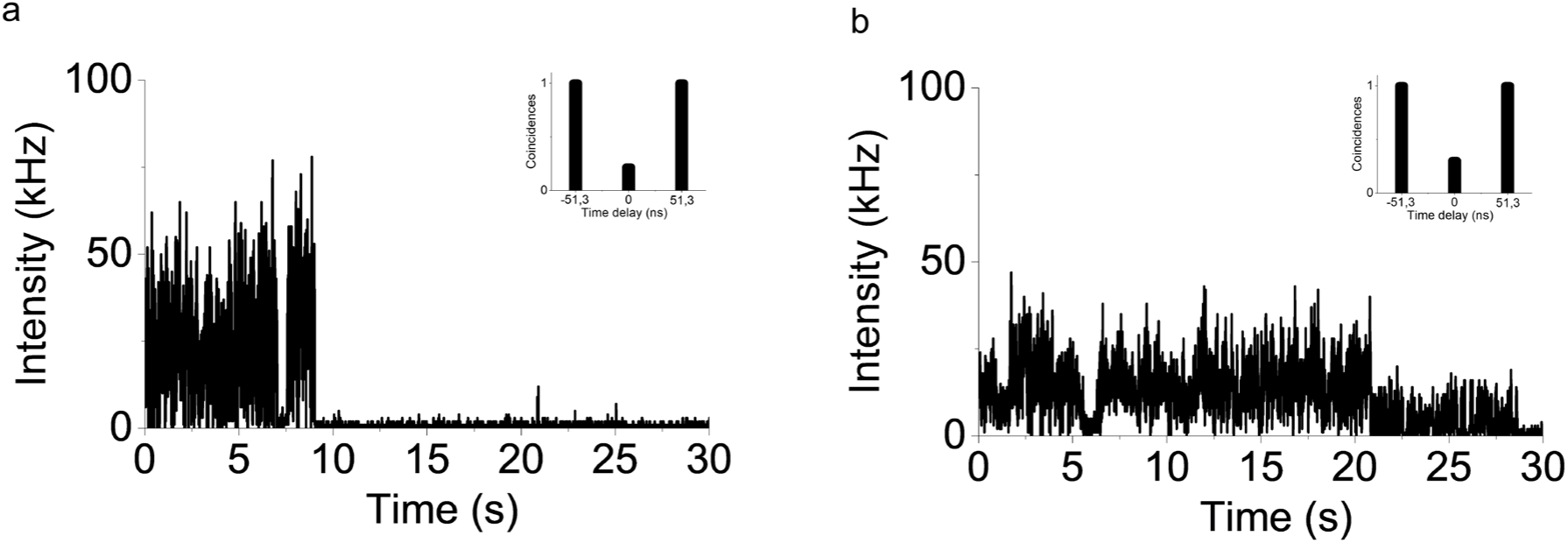
Examples of single-molecule fluorescence trajectories of individual multi-Al647-labeled antibodies in trolox buffer excited at 640 nm with 1 kW cm^-2^ (1 ms binning) that showed signatures of photon antibunching. (**a**) DOL 4.1. (**b**) DOL 8.3. The insets show the normalized *N*_*c*_*/N*_*I,av*_ ratios of the antibunching signal measured for the entire fluorescence intensity trajectories. We determined *N*_*c*_*/N*_*I,av*_ ratios of 0.26 ± 0.02 for the DOL 4.1 antibody (**a**) and 0.31 ± 0.05 for the DOL 8.3 antibody (b).

### Supplementary Videos

**Supplementary Video 1**. Raw *d*STORM data of microtubules in a COS7 cell immunostained with secondary multiple-Al647-labeled antibodies DOL 1.1. *d*STORM movie was recorded using circular polarized laser excitation at 641 nm with an irradiation intensity of 2 kW cm^-2^. Detection was performed at a frame rate of 125 Hz. Buffer conditions: 100 mM MEA with oxygen scavenger, pH 7.6. Scale bars, 1 µm.

**Supplementary Video 2**. Raw *d*STORM data of microtubules in a COS7 cell immunostained with secondary multiple-Al647-labeled antibodies DOL 2.1. *d*STORM movie was recorded using circular polarized laser excitation at 641 nm with an irradiation intensity of 2 kW cm^-2^. Detection was performed at a frame rate of 125 Hz. Buffer conditions: 100 mM MEA with oxygen scavenger, pH 7.6. Scale bars, 1 µm.

**Supplementary Video 3**. Raw *d*STORM data of microtubules in a COS7 cell immunostained with secondary multiple-Al647-labeled antibodies DOL 4.1. *d*STORM movie was recorded using circular polarized laser excitation at 641 nm with an irradiation intensity of 2 kW cm^-2^. Detection was performed at a frame rate of 125 Hz. Buffer conditions: 100 mM MEA with oxygen scavenger, pH 7.6. Scale bars, 1 µm.

**Supplementary Video 4**. Raw *d*STORM data of microtubules in a COS7 cell immunostained with secondary multiple-Al647-labeled antibodies DOL 8.3. *d*STORM movie was recorded using circular polarized laser excitation at 641 nm with an irradiation intensity of 2 kW cm^-2^. Detection was performed at a frame rate of 125 Hz. Buffer conditions: 100 mM MEA with oxygen scavenger, pH 7.6. Scale bars, 1 µm.

## Notes

### Competing Interest Statement

The authors have declared no competing interest.

